# Ligand-induced transmembrane conformational coupling in monomeric EGFR

**DOI:** 10.1101/2021.10.28.466294

**Authors:** Shwetha Srinivasan, Raju Regmi, Xingcheng Lin, Courtney A. Dreyer, Xuyan Chen, Steven D. Quinn, Wei He, Kermit L. Carraway, Matthew A. Coleman, Bin Zhang, Gabriela S. Schlau-Cohen

**Author notes:** These authors contributed equally to this work (Shwetha Srinivasan, Raju Regmi). Department of Physics, University of York, York, UK.

## Abstract

Single pass cell surface receptors regulate cellular processes by transmitting ligand-encoded signals across the plasma membrane via changes to their extracellular and intracellular conformations. While receptor-receptor interactions are established as key aspects of transmembrane signaling, the contribution from the single helix of a monomeric receptor has been challenging to isolate due to the complexity and ligand-dependence of the receptor-receptor interactions. By combining membrane nanodiscs produced wtih cell-free expression, single-molecule Förster Resonance Energy Transfer measurements, and molecular dynamics simulations, we report that ligand binding induces intracellular conformational changes within monomeric, full-length epidermal growth factor receptor (EGFR). Our observations establish the existence of extracellular/intracellular conformational coupling within a single receptor molecule. We implicate a series of electrostatic interactions in the conformational coupling and find the coupling is inhibited by targeted therapeutics and mutations that also inhibit phosphorylation in cells. Collectively, these results introduce a facile mechanism to link the extracellular and intracellular regions through the single transmembrane helix of monomeric EGFR, and raise the possibility that intramolecular transmembrane conformational changes are common to single-pass membrane proteins.

---

Receptor tyrosine kinases, surface receptors present in all cell types across the animal kingdom, regulate major cellular functions, including cell division and survival.^1–3^ Their signaling pathways are initiated by ligand binding to the extracellular region, which causes intracellular autophosphorylation and subsequent recruitment of adaptor proteins to the phosphorylated residues.^1^ Epidermal growth factor receptor (EGFR), a prototypical receptor tyrosine kinase, has been particularly well studied as its aberrant expression leads to diseases such as cancer and diabetes.^4, 5^ Binding of the ligand epidermal growth factor (EGF) induces a conformational expansion of the extracellular region, enabling dimerization of EGFR.^6, 7^ This expansion as well as other ligand-induced changes have been well-characterized for the extracellular region.^8, 9^ The corresponding changes to the intracellular region, however, have only been accessible for oligomers due to the limited window between ligand binding and dimerization.^10^ Analysis of fragmented domains has emerged as an alternative strategy that can isolate the conformations associated with signaling states, yet these domains cannot be used to visualize how extracellular stimuli are propagated across the plasma membrane such as through extracellular/intracellular conformational coupling.^11^ In 47% of membrane proteins, including EGFR, a single trans-membrane helix spans the plasma membrane.^12^ Although different conformations of this helix alone have been observed,^13–15^ how, or even whether, the single helix can support extracellular/intracellular conformational coupling to mediate a signaling cascade has been largely unexplored.

While dimers of EGFR have been long established as an active form of the receptors, emerging evidence suggests that other oligomerization states also play a role in phosphorylation and signaling.^6, 16^ Depending on the lipid composition of the plasma membrane, ligand-induced formation of multimers induces stronger and more complete phosphorylation of the tyrosines and a wider dynamic range of EGFR responsiveness.^16–18^ Furthermore, early studies suggested EGFR signaling can occur even in the presence of an antibody that prevented dimer and multimer formation.^19, 20^ Consistently, certain ligands, such as epigen, lead to monomers or weakly bound dimers, yet still regulate cellular signaling.^21, 22^ Despite these multiple lines of evidence, the behavior of monomeric EGFR and its contribution, if any, to the signaling pathway have not yet been determined. Hence, we used a multidisciplinary approach involving mutagenesis, singlemolecule Förster Resonance Energy Transfer (smFRET), molecular dynamics (MD) simulations, and cellular phosphorylation studies to isolate and investigate extracellular/intracellular conformational coupling within monomeric EGFR and its impact on cell signaling.

## Labeled EGFR monomers in nanodiscs

Cell-free expression was used to produce fulllength EGFR monomers embedded in lipid bilayer nanodiscs and free of cellular interaction partners (Fig. 1a, Supplementary Fig. 1-3).^23^ A FRET donor dye (snap surface 594) was covalently attached to the C-terminus of the protein and an acceptor dye (Cy5) was introduced as a labelled lipid within the bilayer (Supplementary Fig. 4).^24^ The functional, folded conformation of the labelled receptors was implied with ATP-dependent phosphorylation for the intracellular region (Supplementary Fig. 2, Supplementary Table 1, 2) and specificity of ligand binding for the extracellular region (Fig. 2), consistent with previously published western blot and fluorescence-based phosphorylation assays with similar preparations.^23, 24^ Intact, full-length monomeric EGFR was further purified spectroscopically by immobilizing nanodiscs on a cover slip at dilute concentration and only selecting receptors with a single donor and acceptor for analysis (Fig. 1b, Supplementary Fig. 5).

**Figure 1.**
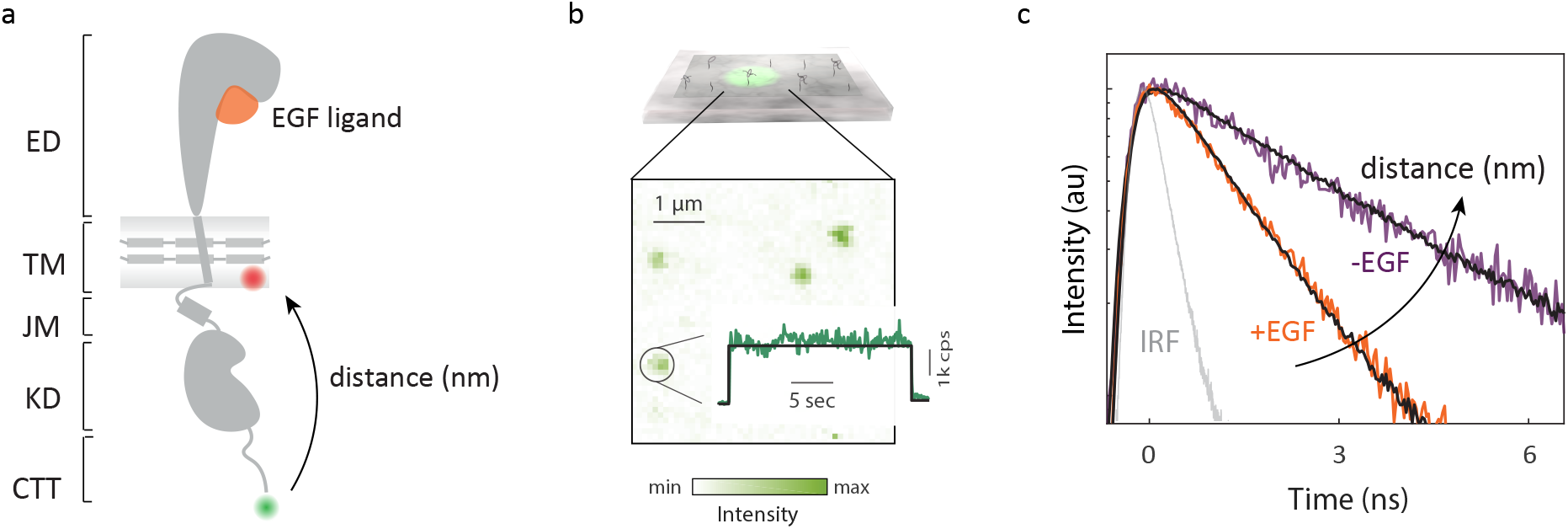
smFRET measures intracellular conformational states of full-length EGFR in a nanodisc. (a) Full-length, monomeric EGFR (solid gray) embedded in a nanodisc. The nanodisc is a lipid bilayer (shaded gray) belted by an amphiphilic apolipoprotein (solid gray). EGFR consists of a 621-amino acid extracellular region (ED) that binds EGF (orange), a 24-amino acid transmembrane-spanning domain (TM), and an intracellular region, which is a 37-amino acid juxtamembrane domain (JM), a 273-amino acid kinase domain (KD) and a 231-amino acid disordered C-terminal tail (CTT) (Supplementary Fig. 1). Green and red spheres indicate the donor and acceptor dyes, respectively. (b) Top: Ni-NTA coated coverslip binds EGFR nanodiscs via a His-tag on the apolipoprotein. Bottom: fluorescence intensity from a confocal image for a representative region *(λ_exc_* = 550 nm) where green spots are immobilized EGFR nanodiscs. Number of detected photons for each 100 ms interval generates a fluorescence intensity trace (green) with the average indicated (black). (c) Histogram of the arrival times of detected photons generates the donor lifetime decay profile. Representative decay profiles of EGFR in the presence (orange) and absence (purple) of the EGF ligand in neutral lipids with the instrument response function (IRF; gray).

**Figure 2.**
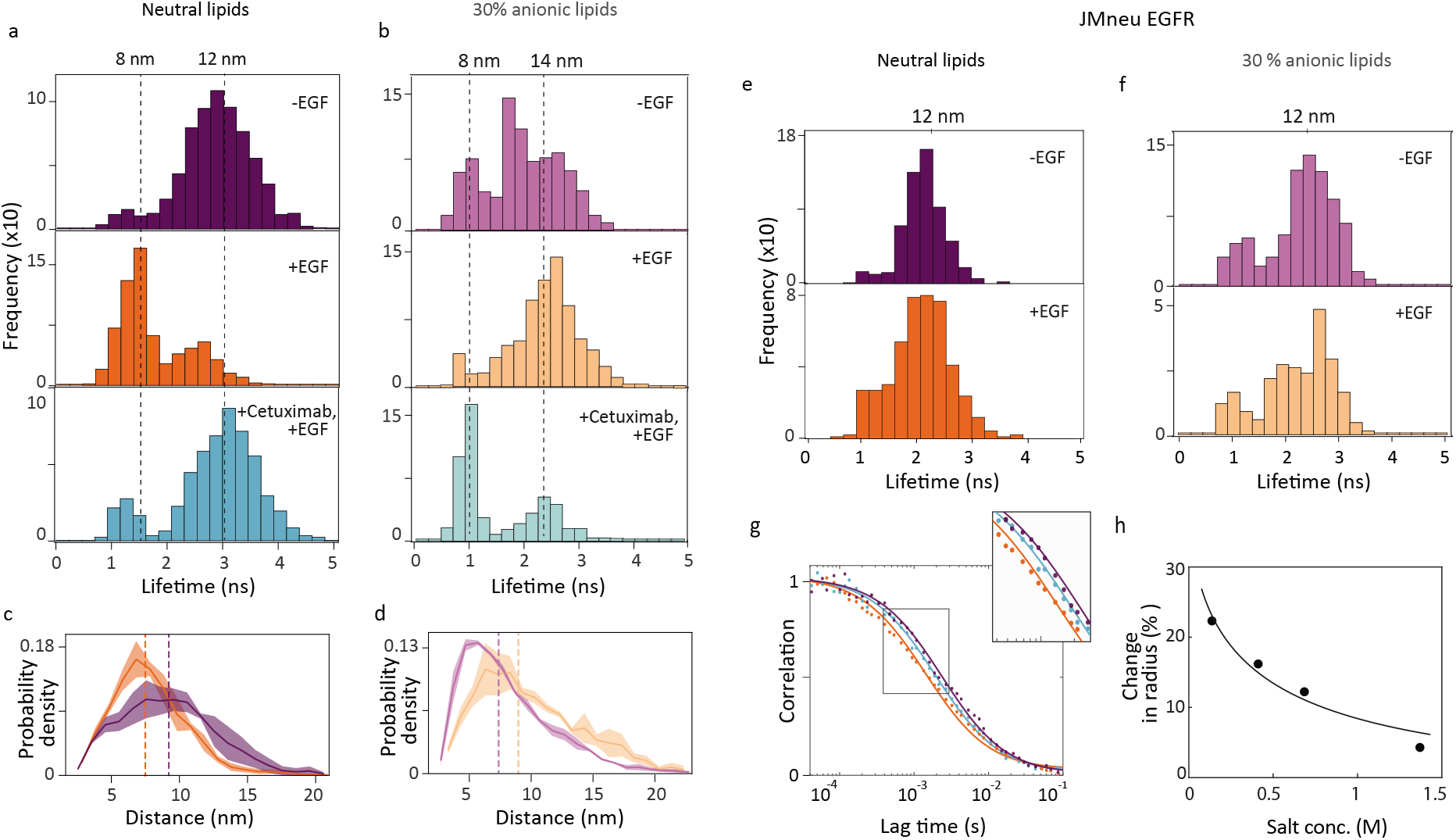
Charged residues implicated in extracellular/intracellular conformational coupling. (a-b): sm-FRET measurements were used to build histograms of the donor lifetime, which shortens as the donor-acceptor distance decreases. Dotted lines indicate the medians with corresponding distances on upper x-axis. The donor fluorescence lifetime distribution in (a) DMPC and (b) POPC-POPS lipids without EGF (top); with 1 *μ*M EGF (middle); with 1 *μ*M EGF and 100 nM Cetuximab. (c-d) Molecular dynamics trajectories were used to find the probability density distribution of the vertical separation in (c) DMPC and (d) POPC-POPS lipids without EGF (purple) and with EGF (orange). Dotted lines indicate the median of the distributions. (e-f) smFRET donor fluorescence lifetime histograms for JMneu EGFR in (e) DMPC and (f) POPC-POPS lipids. Both sets of histograms are statistically similar in the absence (top) and presence (bottom) of EGF (Supplementary Fig. 13). (g) FCS curves of WT EGFR (dots) were fit (solid lines) to extract a diffusion time of 1.5 ms in the presence of EGF (orange), 2.2 ms in the absence of EGF (purple), corresponding to a reduction of 22% in hydrodynamic radius, and 1.9 ms in the presence of Cetuximab together with EGF (cyan). (h) Percent change in hydrodynamic radius from FCS curves in the presence of EGF relative to in the absence of EGF (dots) as a function of salt concentration with fit curve (solid line).

## Intracellular conformational change

The intracellular conformation was examined with smFRET by measuring the fluorescence lifetime of the donor. FRET, which depends on the distance between the donor and acceptor dyes, competes with emission, thereby shortening the lifetime in distance-dependent manner.^25^ The fluorescence lifetime of the donor was measured in the absence and presence of saturating (1 *μ*M) EGF ligand. For monomeric wild type (WT) EGFR embedded in a nanodisc with a 1,2-Dimyristoyl-sn-glycero-3-phosphocholine (DMPC) bilayer, a shorter (2 ×) fluorescence lifetime of the donor was observed in the presence of EGF compared to in its absence for most receptors, signifying a decrease in distance between the C-terminus and the membrane surface upon EGF binding (Fig. 1c).

We built donor lifetime histograms for WT EGFR with and without EGF ligand (Figure 2a). The corresponding donor-acceptor distances were calculated from the lifetimes with reference time *t_D_* = 3.32 ns (Supplementary Fig. 5c). In absence of EGF, the distribution peaked at ~3 ns (12 nm), whereas in the presence of EGF, the distribution peaked at ~1.5 ns (8 nm; Supplementary Figs. 6-8 and Supplementary Table 3). A structural model of EGFR dimers suggests that the kinase domain (KD) of one monomer is lifted towards the membrane, similar to the change seen in the smFRET measurements of monomers.^10^

Fluorescence correlation spectroscopy (FCS) was performed on diffusing donor-only constructs of the full-length, nanodisc-embedded EGFR to characterize the overall structural change of the receptor embedded nanodisc (Fig. 2g; also see Methods). The diffusion coefficient, which is a function of the hydrodynamic radius, can be extracted from transit times through a known confocal volume.^24^ Diffusion coefficients of (1.27 ± 0.08) × 10^-7^ cm^2^ s^-1^ and (0.98 ± 0.03) × 10^-7^ cm^2^ s^-1^ were found in the presence and absence of EGF, respectively, corresponding to a compaction of the hydrodynamic radius (~22%) upon EGF binding (Fig. 2g). Previous crystallographic studies of EGFR reported an expansion in the extracellular region upon EGF binding.^26^ Therefore, the observed compaction likely originates in the intracellular region, consistent with the smFRET results shown in Figure 2a.

We also investigated the effect of Cetuximab, an EGFR antibody administered for metastatic colorectal cancer that inhibits ligand-induced phosphorylation and signaling.^27^ Cetuximab binds at the same extracellular site as EGF but does not cause an extracellular expansion.^28^ In the presence of 100 nM Cetuximab together with 1 *μ*M EGF (Fig. 2a, bottom), the lifetime distributions peaked ~3 ns (12 nm), similar to the distribution in the absence of EGF (Supplementary Figs. 6-8). In FCS measurements, addition of the Cetuximab with EGF produced a diffusion coefficient of (1.10 ± 0.03) × 10^-7^ cm^2^ s^-1^ (Fig. 2g, cyan), which approaches the value in the absence of EGF. The correlation between the extracellular expansion and intracellular compaction indicates the presence of extracellular/intracellular conformational coupling.

While the neutral membrane provided the simplest platform for experiments, we then ascertained the effect of a near-native lipid composition on the intracellular region by incorporating a partially anionic bilayer into the nanodiscs (70% 1-palmitoyl-2-oleoyl-sn-glycero-3-phosphocholine, POPC; 30% 1-palmitoyl-2-oleoyl-sn-glycero-3-phospho-L-serine, POPS; Supplementary Fig. 3), which replicates the plasma membrane anionic lipid content of mammalian cells.^29^ As shown in Figure 2b, smFRET measurements were performed on the receptors in the partially anionic lipid bilayer for all three ligand conditions. In the absence of EGF, the distribution was broad and structured with a median at ~2 ns (10 nm), whereas in the presence of EGF, the distribution appeared closer to unimodal and peaked at ~2.50 ns (≥13 nm; Supplementary Fig. 9 and Supplementary Table 3). Although an EGF-induced intracellular conformational change was again observed, these distributions showed an expansion of the intracellular region, in contrast to the results with the neutral bilayer. In the presence of Cetuximab together with EGF, the distribution peaked at ~1.1 ns (8 nm), which indicated a more compact intracellular region, although one that was again closer to the conformation in the absence of EGF than in its presence. The additional compaction induced by Cetuximab may interfere with the access of substrates and/or signaling proteins to the intracellular binding sites.

## Effect of electrostatic interactions

Given the pronounced effect of anionic lipids on the nature of ligand-induced intracellular conformational changes, we used mutagenesis to investigate the role of electrostatic interactions in extracellular/intracellular conformational coupling. First, we used all atom MD simulations with the CHARMM force field to implicate two regions in the intracellular conformational change (Supplementary Fig. 10), the juxtamembrane domain (JMD), which is the membrane adjacent region of the intracellular domain, and the C-terminal tail (CTT), which contains the tyrosine phosphorylation sites. Next, we exchanged charged residues with their neutral analogs for these regions. To examine the contribution of each domain individually, six of the positively charged residues in the JMD were neutralized in one mutant (JMneu EGFR) and five of the negatively charged residues in the CTT were neutralized in a second (CTTneu EGFR; see Methods, Supplementary Fig. 11).

We performed smFRET measurements on both mutants in neutral and partially anionic lipid bilayers and in the presence and absence of EGF. We built donor lifetime histograms for all eight samples (Fig. 2e and f and Supplementary Fig. 12). In contrast to the distributions for WT EGFR (Fig. 2a, b), the distributions for both mutant samples were statistically similar in the absence and presence of EGF (Supplementary Fig. 13, 14), indicating that the EGF-induced intracellular conformational change is suppressed upon mutation. For three of the samples, JM-neu EGFR in both lipid bilayers and CTTneu EGFR in a neutral bilayer, the distributions peaked at ~12 nm, which is the same distance as for WT EGFR in a neutral bilayer in the absence of EGF; this likely reflects a similar intracellular conformation due to loss of ligand-dependent electrostatic interactions. For CTTneu EGFR in a partially anionic bilayer, the distributions peaked at ~1.2 ns (8 nm), where the decreased distance may be attributed to a loss of repulsion between the negative residues on the tail and anionic bilayer. In contrast to what we observe for WT EGFR, previous structural studies did not show an intracellular conformational change upon EGF binding.^30^ However, the EGFR construct employed in these experiments lacked the CTT, and thus may be more analogous to the measurements of CTTneu EGFR, where an EGF-induced conformational change is similarly not observed.

As additional characterization of the role of electrostatic interactions in the conformations of WT EGFR, we used FCS to measure the EGF-induced compaction of the hydrodynamic radius in a neutral bilayer as a function of salt concentration (Fig. 2h). We observed that the 22% compaction at physiological salt concentration (137 mM) reduced to 4% at high salt concentration (1.37 M). The data points as a function of salt concentration were fit using Debye-Huckel theory for electrostatic screening,^31^ which gave good agreement with the measured values. These results support a model in which extracellular/intracellular conformational coupling is driven by electrostatic interactions.

## Simulations capture measured distances

To further investigate the mechanism behind the observed extracellular/intracellular conformational coupling, we performed explicit solvent MD simulations on full-length, monomeric EGFR. Simulations were carried out on active (+EGF) or inactive (-EGF) conformations in neutral (DMPC) or partially anionic (70% POPC/30% POPS) lipid bilayers for a total of 400 *μ*s using a calibrated Martini force field with improved accuracy in modeling the disordered CTT (Methods, Supplementary Fig. 15).^32^

The average vertical separations between the center of mass of the membrane and the C-terminus (residue 1186) were used to compare the simulated structures with the smFRET results (Fig. 3a). In the neutral bilayer, the average vertical separation stabilized at 9.2 nm and 7.9 nm for the simulated conformations in the absence and presence of EGF, respectively (Fig. 2c). In contrast, in the partially anionic bilayer, the average vertical separation was longer in the presence of EGF (9.8 nm) than in its absence (8.1 nm; Fig. 2d). Therefore, comparison amongst the intracellular domains of the simulated structures showed that the simulations succeeded in reproducing the experimental trends (Fig. 2c, d, Supplementary Fig. 16).

**Figure 3.**
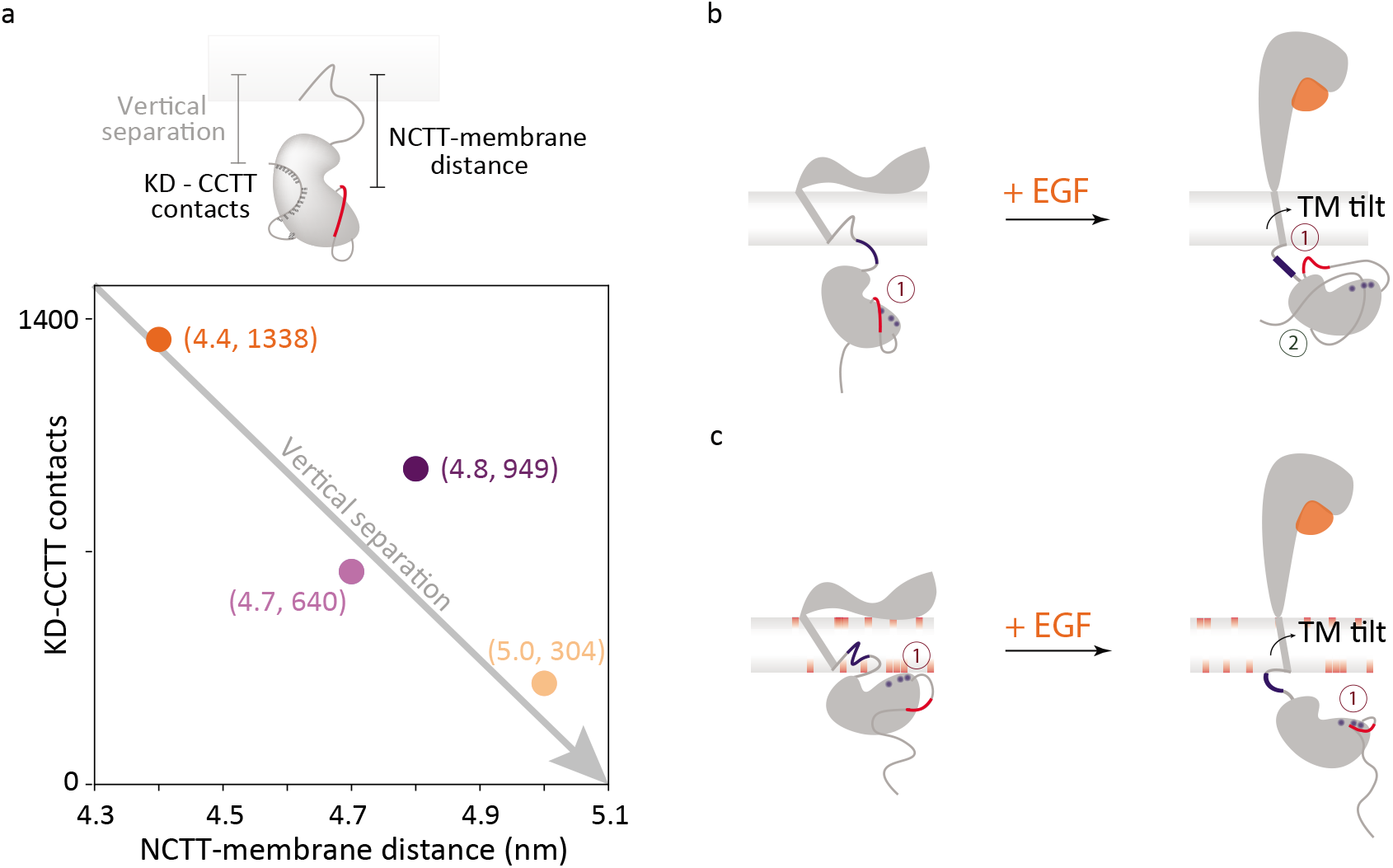
Intracellular conformational changes upon extracellular EGF binding. (a) The intracellular conformation can be characterized by the distance between the NCTT and the membrane and the contacts between the CCTT and the KD. The positions of WT EGFR in both neutral (dark colors) and partially anionic (light colors) lipid bilayers in the absence (purple) and presence (orange) of EGF are plotted as a function of the average NCTT-membrane distance and KD-CCTT contacts obtained from simulations. (b-c) Schematic of proposed mechanism for EGFR (gray) conformational changes upon EGF (orange) binding in (b) neutral and (c) partially anionic membrane. Numbers indicate electrostatic (dark red) and hydrophobic (dark green) interactions. (b) Left: the flat extracellular domain tilts the transmembrane helix, pulling the JMD (blue line) into the lipid bilayer. The negatively charged NCTT (red) interacts with the positive residues in the KD (blue) leading to the release of the CCTT. Right: with EGF bound, the upright extracellular domain allows the transmembrane helix to be vertical, causing the full JMD to protrude from the membrane. The positively-charged JMD (blue) attracts (1) the negatively-charged CTT (red). In combination with hydrophobic interactions between the CCTT and KD (2), the attraction produces an intracellular compaction. (c) Left: the transmembrane tilt and the anionic lipids embed the positively-charged JMD into the membrane, which pulls the positively-charged KD (blue) closer to (1) interact with the negatively-charged lipids (red). Right: with EGF bound, the upright transmembrane domain extends the JMD out of the membrane, further positioning the KD away from the membrane. The positive residues in the KD (blue) interact (1) with the negatively-charged CTT (red).

## Transmembrane conformational coupling

Collectively, the simulations indicate that the overall intracellular conformation can be characterized based on two parameters: the distance between the N-terminal portion of the CTT (NCTT) and the plasma membrane and the number of contacts between the C-terminal portion of the CTT (CCTT) and the KD (Supplementary Fig. 17), as illustrated in Fig. 3a. For the structures that corresponded to the smFRET measurements, both of these parameters were strongly correlated with the measured distances (Supplementary Fig. 17-19). Examination of the simulations, along with the experimental data and previously published results, points to a molecular mechanism for extracellular/intracellular conformational coupling, as illustrated in Figs. 3b and c.

With EGF bound, the extracellular domain extends above the membrane,^33^ imposing a vertical orientation to the transmembrane domain (TMD) (Supplementary Fig. 20). Without EGF, the hinge action of the extracellular domain causes it to lay flat on the membrane,^33^ imposing a tilted orientation to the TMD (Supplementary Fig. 20). The tilted TMD positions the JMD closer to the membrane surface, which increases the JMD–lipid interactions. The position and interactions of the JMD in turn influence the conformation of the entire intracellular region of the receptor.

In the neutral bilayer (Fig. 3b), EGF binding exposes the positively-charged residues of the JMD for electrostatic interactions, including with the negatively-charged residues of the CTT, leading to a more compact structure. In both JMneu and CTTneu EGFR, this interaction —and thus the EGF-dependent conformational change —are absent (Fig. 2e, Supplementary Fig. 12). In the partially anionic bilayer (Fig. 3c), the positively charged residues of the JMD tend to embed into lipid bilayer (Supplementary Fig. 21), lifting the KD and thus the intracellular domain closer to the membrane (Supplementary Fig. 18–19). In this case, the decreased tilt of the TMD with EGF bound extends the JMD, positioning the KD and subsequently the CTT further from the plasma membrane. The neutralization in JMneu EGFR likely frees the JMD from the anionic membrane, leading to an intracellular expansion similar to the case of the neutral bilayer without EGF (Fig. 2f). The neutralization in CTTneu EGFR, on the other hand, removes the repulsion between the CTT and the anionic membrane, leading to a more compact intracellular domain that still lacks the EGF-dependent effects observed in WT EGFR (Supplementary Fig. 12).

## Phosphorylation reduced by neutralization

To explore the consequences of the intra-cellular compaction in signaling, phosphorylation was monitored for wild-type and mutated EGFR in CHO cells in the absence and presence of EGF (Fig. 4a). JMneu EGFR showed reduced phosphorylation by 40%-60% when compared to WT EGFR (Fig. 4b; Supplementary Table 4; Supplementary Fig. 22). Consistent with these observations, in previous studies it was found that deletion of the charged segment of the JMD reduced phosphorylation levels by 95% and neutralization of individual residues reduced levels by up to 50%.^34–36^ In addition, substitution of the native sequence with a neutral, unstructured sequence led to the disappearance of signaling, even while ligand binding and dimerization capabilities were retained.^37^ The JMD has been shown to mediate autophosphorylation efficiency *via* conformational changes that differ with ligand identity,^38^ possibly reflecting the changes we observe here (Fig. 4c). While both reduction of phosphorylation levels in cells and a loss of extracellular/intracellular conforma-tional coupling in vitro were seen for JMneu EGFR, interpretation of the cellular phosphorylation results is complicated by the presence of interactions with other receptors or charged proteins.^33, 36, 39^

**Figure 4.**
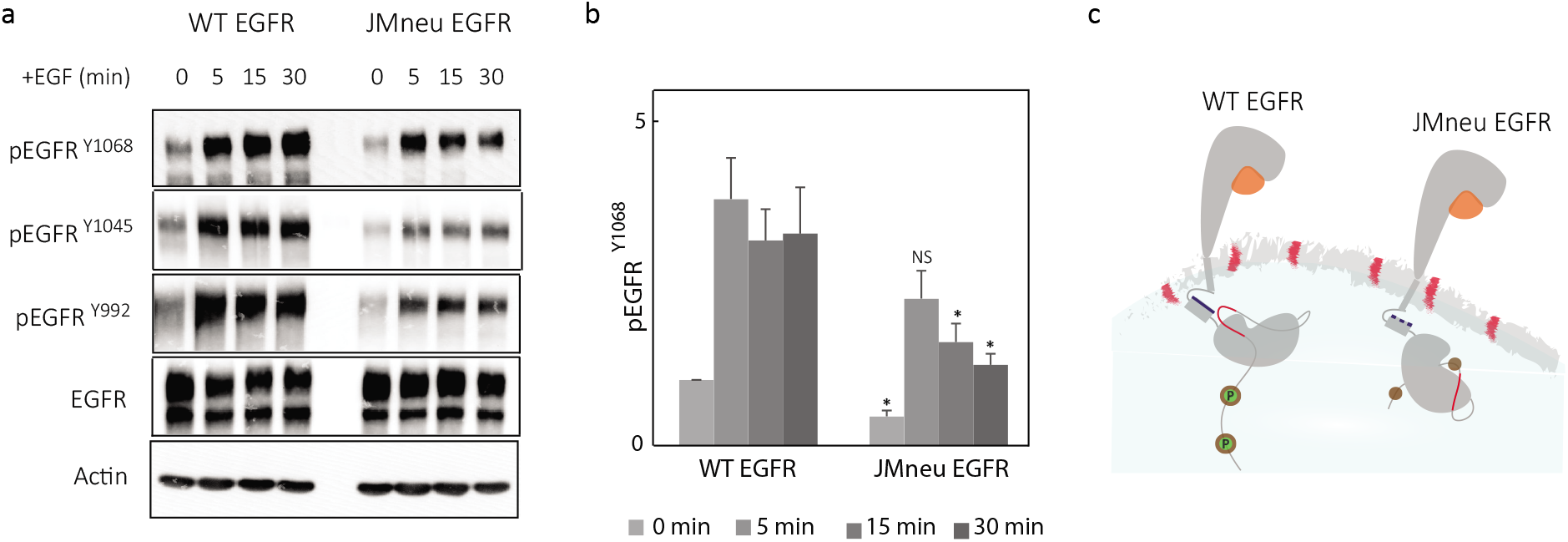
Cellular experiments with JMneu EGFR show reduced phosphorylation. (a) Expression of phosphorylated EGFR (pEGFRY1068, pEGFRY1045, pEGFRY992) and total EGFR at 0, 5, 15, and 30 minutes of 100 ng/mL EGF stimulation in CHO cells transfected with WT EGFR and JMneu EGFR. Actin was used as a loading control. (b) Relative expression of pEGFRY1068 was determined by western blot quantification. The values were normalized to pEGFRY1068 before EGF stimulation (time 0) of the wild-type EGFR transfected cells. The results reported are representative of six independent biological replicates; *p<0.05. (c) Schematic of proposed effect of JMneu EGFR on phosphorylation in CHO cells. The positive region in the JMD is shown in blue in WT EGFR and in dotted blue in JMneu EGFR. The negative lipids and region in the NCTT are shown in red. The circles in the CTT indicate tyrosine phosphorylation (P) sites.

Phosphorylation was also monitored for the CTTneu EGFR (Supplementary Fig. 23), where a high basal level of phosphorylation was observed due to the previously established autoinhibitory role of the mutated residues.^40^ However, only a marginal increase in phosphorylation was measured upon addition of EGF, demonstrating that ligand-dependent phosphorylation also decreases relative to WT EGFR upon neutralization of the CTT. Dimerization of wild-type EGFR in the absence of ligand binding was previously shown to be insufficient for signaling, and was ascribed to an unidentified EGF-induced conformational change, such as the one identified in this study.^41, 42^

## Discussion

Collectively, our observations indicate that conformational changes within the intracellular region of the EGFR are dictated by the previously-reported ligand-induced conformational changes in the extracellular region. We envision that the ligand-induced and lipid-dependent conformations within the monomer precede ligand-induced dimerization. The correlation between the extracellular/intracellular conformational coupling and the phosphorylation levels in cells suggests that the coupling may be a part of the biophysical mechanism of signaling. The variable intracellular conformation may also contribute to the differences in signaling efficiency and autophosphorylation site usage induced by different ligands.^43, 44^

Electrostatic interactions appear to define the intracellular conformation of a receptor monomer, with the positively-charged JMD as a key mediator, and so lipid charge has a profound impact on the nature of the conformations. Our results complement and expand upon previously identified electrostatic interactions between anionic lipids and the positively-charged regions of the JMD and KD, which were reported to restrict access of the substrate tyrosines to the KD and thereby regulate phosphorylation.^33, 36, 38, 45^ Similarly, ligand-induced binding of a calcium/calmodulin complex to the JMD was found to reverse the local net charge and detach the KD from the membrane, exposing the ATP binding region for substrates.^39, 46^

The studies presented here demonstrate that extracellular/intracellular conformational coupling is achieved through the single transmembrane helix of monomeric EGFR. In the seven transmembrane helices of rhodopsin and several other G-protein coupled receptors, highly conserved charged residues encompassing a network of electrostatic interactions are thought to lock the protein in its inactive conformation, regulating the visual transduction signalling cascade.^47^ Similarly, multipass proteins have also been shown to transition into an active form through small orientational changes in the transmembrane region upon ligand binding.^48, 49^ Our results suggest a similar effect can be achieved within a single pass protein where one *α*-helix serves as a minimal yet sufficient system for signal transduction. This system may be shared by other single pass membrane proteins with similar structures and functions.^50^

## Acknowledgments

This work was supported by the NIH Directors New Innovator Award 1DP2GM128200-01 (to G.S.S.-C.). Research was also supported by the Laser Biomedical Research Center (NIH9P41EB015871). X.L. and B.Z. acknowledge support by startup funds from the Department of Chemistry at the Massachusetts Institute of Technology. S.D.Q. acknowledges support from the Lindemann Trust. This work was in part performed under the auspices of the U.S. Department of Energy by Lawrence Livermore National Laboratory under Contract DE-AC52-07NA27344. Research was supported in part by the National Institutes of Health under award numbers R21AI120925, R01CA155642 and R01GM117342 (to M.A.C.).

## Author contributions

S.S., R.R., K.L.C., M.A.C., and G.S.S.-C conceived the experiments. W.H., K.L.C, and M.A.C. prepared the EGFR and ApoA1 plasmids. S.D.Q., S.S., and R.R. optimized labeled EGFR production. S.S. and R.R. performed and analyzed the fluorescence experiments. X.L. and B.Z. designed and performed the simulations. C.A.D. performed the cell culture experiments. X.C. performed the EGFR function-related and characterization experiments. S.S., R.R., and G.S.S.-C. co-wrote the manuscript. All authors discussed the results and commented on the manuscript.

## Supplementary Information

Supplementary Information is available for this paper.

## Data availability statement

The data that support the findings of the present study are available from the corresponding authors upon request.

## Corresponding authors

Correspondence to Bin Zhang and Gabriela S. Schlau-Cohen.

