## Supplementary Information for "Ligand-induced transmembrane conformational coupling in monomeric EGFR"


This document contains the following supporting information:


Methods

Figures S1 – S24

Tables S1 – S9

References (1 - 28)

### Methods

#### Production of labeled full-length EGFR nanodiscs

Fluorescently-labelled EGFR in nanodiscs was produced and characterized by adapting previous published protocols.[1,2] Plasmid encoding 6× His-tagged version of human apolipoprotein A1 lacking the amino-terminal (ApoA1Δ49) and full-length EGFR (1210 amino acids) with a SNAP tag at the C-terminal position were codon optimized for $Escherichia\ Coli$ ($E.\ Coli$) expression in pIVEX2.4d and SNAP-T7 vector respectively (purchased from GenScript). Cell-free expression was carried out utilizing the Expressway Maxi Cell-Free $E.\ Coli$ Expression system (Life Technologies) to produce full-length EGFR fused with a SNAP tag (Supplementary Fig. 2a). A final concentration of 2 mg/mL of probe sonicated 1,2-dimyristoyl-sn-glycero-3-phosphocholine (DMPC) lipid (Avanti Polar Lipids) vesicles in distilled water was added to the $E.\ Coli$ Sly D extract, *in vitro* protein synthesis (IVPS) $E.\ Coli$ reaction buffer, amino acids (without methoninine), methionine, T7 enzyme mix, DNA templates to make a neutral synthetic bilayer. To mimic a partially anionic symmetric bilayer, 2 mg/mL of 70% 1-palmitoyl-2-oleoyl-sn-glycero-3-phosphocholine (POPC) (Avanti Polar Lipids) probe sonicated lipid vesicles and 30% 1-palmitoyl-2-oleoyl-sn-glycero-3-phospho-L-serine (POPS) (Avanti Polar Lipids) probe sonicated lipid vesicles were used in the cell-free reaction. A molar ratio of 500:1 lipid:Cy5 labeled 1,2-dioleoyl-sn-glycero-3-phosphoethanolamine (DOPE) lipids (Avanti Polar lipids) was added to the above solutions after bath sonication to introduce a single acceptor in the nanodisc. The lipid molar ratio was empirically optimized for one acceptor per nanodisc (Supplementary Fig. 4). 20 $\mu$g of EGFR DNA and 0.2 $\mu$g of ApoA1Δ49 DNA were added to the lysate in addition to protease inhibitor cocktail (ThermoFisher Scientific) and RNAse inhibitor (Roche). The solution was incubated at 200 rpm for 30 minutes at 25$^o$ C. A feed buffer solution was made using IVPS solution, amino acids (without methionine) and methionine. A total reaction volume of 250 $\mu$L was incubated for a total time of 10 hours at 200 rpm at 25$^o$ C. Prior to the protein purification, 500 nM of snap surface 594 (New England Biolabs) was added to the reaction and was incubated at 37$^o$ C, 150 rpm for 35 minutes for fluorescence labeling. Snap surface 594 is a derivative of Atto 594 with benzylguanine functionality that reacts with the genetically-encoded snap tag with near-stoichiometric efficiency.[3] For experiments measuring the distances between residues 721 and C-terminus of the protein, atto647N labeled gamma-ATP (Jena Bioscience) was added along with snap surface 594.[2]

For experiments with a neutralized juxtamembrane-A (JMA) domain, JMneu EGFR was made with positively charged residues Arg651, Lys652, Arg653, Arg656, Arg657 and Arg662 in WT EGFR mutated to neutral alanine residues. The mutations in the JMA domain are unlikely to affect helix formation or hydrophobic interactions with the membrane bilayer as the hydrophobic residues of JMA were undisturbed. Similarly, for experiments with a neutralized C-terminal tail (CTT), CTTneu EGFR was made with negatively charged residues Asp984, Asp985, Asp988, Asp990, Glu991 in WT EGFR mutated to neutral residues Asn984, Asn985, Asn988, Asn990 and Gln 991 (see Supplementary Fig 10). The same protocol as above was used for cell free expression of mutated EGFR receptors using plasmids with these modified DNA sequences.

**Affinity Purification of labeled EGFR nanodiscs**

500 $\mu$L of Ni-NTA resin slurry (Qiagen) was added to a 2 mL plastic column (Bio-Rad Laboratories). The resin was washed with double distilled water and equilibrated with 3 mL of native lysis buffer (50 mM $NaH_2PO_4$, 300 mM NaCl, pH 8.0). The cell-free reaction post labeling was added to 500 $\mu$L of lysis buffer on the incubated column and incubated at 4$^o$ C for 2 hours. The flowthrough was collected and the column was washed with lysis buffer (10 $\times$ 1 mL) followed by lysis buffer containing 10 mM imidazole (2 $\times$ 1 mL), lysis buffer containing 25 mM imidazole (2 $\times$ 1 mL), and lysis buffer containing 50 mM imidazole (2 $\times$ 1 mL) to remove all the

non-specific interactions of the reaction mixture and free dye from the column. The EGFR nanodiscs were eluted with lysis buffer containing 400 mM ($2 \times 500$ $\mu$L) imidazole. The samples were finally concentrated using 50 kDa, 500 $\mu$L spin filters (Sigma-Aldrich) by centrifugation.

**Protein content for labeled EGFR nanodiscs**

SDS-PAGE was used to confirm the production of both belt protein (at 25 kDa) and EGFR (at 160 kDa) (Supplementary Fig. 2b). Samples were mixed with 2$\times$ Laemmli sample buffer (Bio-Rad Laboratories), 2.5% 2-mercaptoethanol (Sigma-Aldrich) and boiled for 5 minutes at $100^o$ C before running on precast stain free gels from Bio-Rad Laboratories. Precision Plus Protein unstained standard (Bio-Rad Laboratories) marker was used for the stainfree imaging and Prestained NIR protein ladder (ThermoFisher) for fluorescence imaging. Gels were run at 170 V for 45 minutes. Stain-free imaging was performed on a Gel Doc imager (Bio-Rad Laboratories) and fluorescent images were acquired form Typhoon Gel FLA 9500 imager (GE Healthcare Life Sciences). The other proteins appearing in the stainfree gel are proteins not expressed completely during the cell-free reaction or the transcription and translation machinery of the cell-free reaction mixture. The specificity of snap surface 594 fluorophore binding to EGFR was confirmed through a fluorescence gel (Supplementary Fig. 2b).

**Transmission Electron Microscopy**

5 $\mu$L of cell-free expressed EGFR nanodiscs in 1$\times$ PBS buffer (137 mM NaCl, 2.7 mM KCl, 10 mM $Na_2HPO_4$, 1.8 mM $NaH_2PO_4$, pH 7.4) were added to glow-discharged carbon coated 400 mesh copper grids (Electron Microscopy Sciences) and incubated for 5 minutes at room temperature to allow nonspecific binding of the nanodiscs to the grids. The solutions were removed by gently blotting the side of the grid with filter paper. The grids were subsequently incubated with 5 $\mu$L of 2 % aqueous uranyl acetate for 30 seconds. Excess stain was removed

similarly to the sample. The grids were air dried, and then imaged on a FEI Tecnai transmission electron microscope (120 kV, 0.35 nm point resolution). The distribution of disc sizes was analyzed using Image J software.

#### Dynamic light scattering

The EGFR nanodiscs in $1\times$ PBS buffer were filtered using 0.2 $\mu$m syringe filters and their dynamic light scattering measurements were performed on a DynaPro NanoStar (Wyatt Technologies, USA). Each measurement represents an average of 50 individual runs.

#### Zeta potential measurements to quantify surface charge of nanodiscs

Titrations (0%, 30%) of negatively charged lipid (PS) in neutral lipid (PC) were performed to determine the surface charge of the nanodiscs with increasing negatively charged lipid content. Zeta potential measurements were performed on a Malvern Zetasizer Nano - ZS90 (Malvern, UK), with a backscattering detection at a constant 173 scattering angle, equipped with a laser at 4mW, 633 nm. Dip cell ZEN1002 (Malvern UK) was used in the zeta-potential experiments. EGFR loaded nanodiscs produced from cell-free reactions were purified as mentioned above and buffered exchanged to 0.1 $\times$ PBS. A final volume of 650 uL was prepared and transferred into the zeta dip cell. For each sample, a total of five scans, 30 runs each, with an initial equilibration time of 5 minutes, were recorded. All experiments were performed at 25$^o$ C. Values of the viscosity and refractive index were set at 0.8878 cP and 1.330, respectively. Data analysis was processed using the instrumental Malverns DTS software to obtain the mean zeta-potential value.

#### Phosphorylation of EGFR in nanodiscs

Western Blot was performed using the Trans-Blot Turbo Transfer System from Bio-Rad Laboratories. The pre-loaded program for high molecular weight protein transfer was used for membrane transfer. After the transfer, the membrane was blocked in 5% Non Fat Dry Milk (prepared in TBST buffer) for the anti-EGFR Western blots, and in 5% BSA (prepared in TBST buffer) for the anti-phosphotyrosine Western blots for 20 minutes at room temperature. The membrane was then incubated in primary antibody overnight at $4^o$ C. Following day, the membrane was washed and incubated with secondary antibody for 1 hour at room temperature. The primary antibodies, secondary antibodies and dilutions are listed in Supplementary Table 8. The fluorescence band detection was done using the ChemiDoc Imaging System from Bio-Rad Laboratories.

Phosphorylated tyrosines were also detected using an Y1068 antibody, which binds specifically to the phosphorylated tyrosine pY1068. 1 nM of snap surface 594 labeled EGFR nanodiscs was incubated with 0.25 nM of Y1068 antibody (alexa647 labeled or unlabeled) in the presence of 50 $\mu$M ATP, 15 mM $MnCl_2$, 15 mM $MgCl_2$ and 2 mM DTT for 30 minutes at room temperature. FCS experiments were performed on the mixture as described below. Ensemble FRET measurements were performed by exciting 1 nM of snap surface 594 labeled EGFR nanodiscs in the absence and increasing amounts (0.5 nM, 2 nM, 20 nM) of alexa 647 labeled Y1068 antibody at 550 nm. The time resolved donor fluorescence decay was deconvolved with the instrument response function and fit to a mono-exponential.

**Preparation of ligands (EGF, neuregulin) and Cetuximab drug**

Human EGF produced in $E.\ Coli$ was purchased from Gold Biotechnology (Catalog Number: 1150-04-100). 1 $\mu$M EGF was prepared in 1$\times$ PBS buffer. Cetuximab produced in CHO cells was purchased from Selleckchem (Catalog Number: A2000). 100 nM Cetuximab was prepared in 1$\times$ PBS buffer. Both EGF and Cetuximab were used at concentrations higher than their dissociation constants (1 $\mu$M EGF and 100 nM Cetuximab) to ensure saturation of EGFR molecules.[4,5] Human neuregulin produced in $E. Coli$ was purchased from Cell Signaling Technology (Catalog Number: 5218SF). 1 $\mu$M neuregulin was prepared in 1$\times$ PBS buffer.

**Fluorescence Spectroscopy**

The His-tag present on the belt protein (ApoA1$\Delta$49) was used to immobilize the EGFR-nanodisc constructs onto the microscope coverslip *via* Ni-NTA affinity. The purified EGFR nanodiscs were diltued to $\sim$500 pM in 1$\times$ PBS buffer and incubated for 15 minutes on the Ni-NTA coated glass (from Microsurfaces, Inc.) and flushed with solution containing 2 mM 6-hydroxy-2,5,7,8-tetramethylchroman-2-carboxylic acid (Sigma-Aldrich), 25 nM protocatechuate-3,4-dioxygenase (Sigma-Aldrich) and 2.5 mM protocatechuic acid (Sigma-Aldrich). Fluorescence experiments were then carried out on a home built confocal microscope.[6] A Ti-Sapphire laser (Vitara-S, Coherent: $\lambda_c$ = 800 nm, 70 nm bandwidth, 20 fs pulse duration, 80 MHz repetition rate) was focused into a non-linear photonic crystal fiber (FemtoWhite 800, NKT Photonics) to generate a supercontinuum. Excitation light was then spectrally filtered for pulses centered at 550 nm or 640 nm and focused with an oil immersion objective lens (UPLSAPO100$\times$, Olympus, NA = 1.4). Fluorescence emission was collected by the same objective and fed to the avalanche photodiodes (SPCMAQRH-15, Excelitas). A 5 $\mu$m $\times$ 5 $\mu$m area of a coverslip with immobilized receptors was scanned. Diffraction limited and spatially separated single molecule spots were then probed individually by unblocking the laser beam to record fluorescence until photo-bleaching. For 550 nm excitation, fluorescence was separated with a dichroic filter SP01-561RU (Laser 2000) and passed through FF01-629/56-25 (Semrock) for donor fluorescence collection. For experiments at 640 nm, ET 645/30$\times$ (Chroma) was used as the excitation filter, FF01-629/56-25 (Semrock) as the dichroic and FF02-685/40-25 (Semrock) for acceptor fluorescence collection. The laser power for the experiments was 2-3 $\mu$W at the sample plane.

Florescence emission was binned at 100-ms resolution to generate fluorescence intensity traces for both the donor and acceptor channels. Traces with a single photobleaching step for the donor and acceptor were considered for further analysis. Regions of constant intensity in the traces were identified by a change-point algorithm.[7] Donor traces were assigned as FRET levels until acceptor photobleaching. Consecutive bunches of 1000 photons in the donor channel were used to construct fluorescence decay curves for the FRET levels.[8] The photons were histogrammed and the distributions were fit to a mono-exponential function convolved with the instrument response function (IRF) and summed with a separately-measured background term. The fit was performed using a Maximum Likelihood Estimator (MLE), which has been shown to be more accurate in the single-molecule regime.[6,9] The extracted lifetimes were used to construct histograms with bin sizes estimated from the square root of the total number of photon bunches.

The donor-acceptor distance (r in nm) was estimated using the following relation:[10,11]

$$r = r_o \sqrt[6]{\frac{1-E}{E}}$$

where $r_o$ is the calculated Förster distance (8.49 nm for snap surface 594 & cy5 dye pair)[10] and the FRET efficiency (E) being experimentally measured as:

$$E = 1 - \frac{\tau_{DA}}{\tau_D}$$

$\tau_{DA}$ is the fluorescence lifetime of the donor in the presence of an acceptor, and $\tau_D$ is the lifetime of donor-only construct. The distance between the donor and acceptor was quantified using a reference lifetime determined with a separately-characterized donor-only construct. For the smFRET measurements on the labelled constructs, the Cy5-lipid is confined to one side of the membrane for the duration of the measurement,[12] and so the extracted distances are an average over the rapid translational diffusion of the labelled dye across the surface of the membrane nanodisc.[13]

For fluorescence correlation spectroscopy (FCS) experiments, the same confocal microscope was used to probe diffusing EGFR embedded nanodisc systems at $\sim$1 nM with the focus of the microscope objective adjusted 30 $\mu$m above the coverslip. The temporal fluctuations in the fluorescence signals were autocorrelated to generate the FCS curves, which were then analyzed using a 1-species translational diffusion model:[2]

$$G(\tau) = 1 + \frac{1}{N} \frac{1}{(1 + \tau/\tau_d)\sqrt{1 + s^2\,\tau/\tau_d}}$$

where N is the number of molecules, $\tau_d$ represents the mean residence time (set by translational diffusion) and $s$ being the ratio of transversal to axial dimensions of the confocal volume. The translation diffusion times were then used to extract the diffusion constant D = $\omega^2$/ 4$\tau_d$, where $\omega$ represents the beam waist of the laser focus. The effective hydrodynamics radius (R) was finally derived using Stokes-Einstein relation:

$$D = \frac{k_B T}{6\pi\eta R}$$

where $k_B$ is the Boltzmann's constant, T being the temperature and $\eta$ being the viscosity of the medium.[14] All experiments were performed at room temperature (21$^o$ C) with 30$\%$ humidity. The photon arrival times were recorded by a time-correlated single-photon counting (TCSPC) module (PicoHarp 300, PicoQuant).

The change in radius ($\%$) as a function of salt concentration in Fig. 2h was fit to the below equation following Debye-Huckel theory for electrostatic screening:[15]

$$\Delta R = a e^{-b\sqrt{c}}$$

where $\Delta R$ refers to the change in radius ($\%$) and c is the salt concentration. Fit values are reported in Supplementary Table 5.

#### Statistical information

Statistical analysis was performed using MATLAB. One-way analysis of variance (ANOVA) was performed on different pairs of experimental data and statistical significance was set at P $\leq 0.001$. The P-values, degrees of freedom, F-statistics is reported in Supplementary Table 6. The number of data points in the smFRET lifetime distributions is reported in Supplementary Table 7. The number of data points are from two to four biological replicates for each sample.

#### Coarse-grained, Explicit-solvent Simulations with the MARTINI Force Field

Our MD simulations used the active and inactive structures assembled in a previous work.[16] The active monomer was modeled with one chain of the active dimeric structure, which included only a fraction of the CTT (residue ID: 978-995) solved in the crystal structure (residue ID 1-995). We also constructed a chimera inactive monomer (residue ID 3-990) by replacing the extracellular domain of the active monomer with that of the inactive monomer. Modeller[17] was then used to build the CTT (residue ID: 955 to 1186) as a disordered chain to complete the two protein structures. We carried out a set of simulations of the full-length EGFR, including the disordered CTT, using the coarse-grained MARTINI force field.[18] Though coarse-grained, these simulations provide explicit representations for the lipid bilayer, ions, water molecules, and the protein. They allow direct comparisons with the distances measured in FRET experiments.

Since the Martini force field was originally calibrated for ordered proteins, when directly applied to disordered regions, the simulations produced overly collapsed configurations. Therefore, we adjusted the interaction strength between protein and water molecules to provide more reasonable conformations for the C-terminal domain. Similar modifications have been made to atomistic force fields to improve their accuracy in modeling disordered proteins.[19] Specifically, we scaled the Lennard-Jones potentials between water molecules and all protein atoms of the C-terminal domain (residues 961-1186). The scaling factor, 1.12, was chosen to best reproduce the radius of gyration of the CTT as measured by the SAXS experiment[20] (Supplementary Fig. 15). The interaction between water molecules and other parts of the EGFR was left unchanged. The time step of Martini simulation was set as 20 fs. The simulations were performed at 303K.

With the recalibrated force field, we performed four sets of simulations of the active/inactive EGFR embedded in DMPC/POPC-POPS bilayers, respectively, for a total of 400 $\mu$s. Umbrella sampling[21] was used to facilitate the exploration of the conformational space. The contact number between the KD and the CTT, which captures the interaction strength between these two domains, was selected as one of the collective variables. The contact number was defined as the number of atom pairs between the KD and the CTT that have a smaller than 8 Å mutual distance. The reference umbrella coordinates varied from 1000 to 4000 with a spacing of 750. To accelerate the equilibration of the system toward reference umbrella values, we used a time-dependent spring constant that increased from 0.0 to 0.0005 kcal/mol in the first 10 ns, stayed constant for the following 80 ns, and decreased linearly to 0.000005 kcal/mol in the next 10 ns. The spring constant was kept at 0.000005 kcal/mol for the remaining of the simulations. We performed an additional set of umbrella simulations that included biases over the JMA-NCTT distance, in addition to the contact number between the KD and the CTT, with the target value of 15.0 Å and the spring constant of 0.005 kcal/mol/Å$^2$.

We analyzed the simulation data using WHAM[22] to remove the effect of umbrella biases and compute the true thermodynamic averages of various metrics presented in the main text and supplementary figures 17 - 21, 24.

#### Atomistic Simulations with the CHARMM Force Field

We carried out microseconds long all-atom, explicit solvent simulations to characterize active and inactive EGFR embedded in the DMPC lipid bilayer. Initial configurations of these simulations were constructed using the active and inactive structures assembled in a previous work.[16] The active monomer was modeled with one chain of the active dimeric structure, which included only the fraction of CTT (residue ID: 978-995) solved in the crystal structure (residue ID 1-995). The rest of the CTT was not included in these simulations to reduce system size. We also constructed a chimera inactive monomer (residue ID 3-990) by replacing the extracellular domain of the active monomer with that of the inactive monomer. Both the active and chimera inactive monomers of EGFR were embedded in a neutral DMPC membrane, prepared with the CHARMM-gui toolkit.[23] Each system was simulated in a NPT ensemble for 1.5 $\mu$s with the time step of 2 fs. CHARMM36m force field[24] and TIP3P water molecules[25] were used. The systems were neutralized and solvated with 0.15 M NaCl, reaching 351,139 atoms used for the simulation of active monomer, and 294,387 atoms used for the simulation of chimera inactive structure. The simulations were performed using GROMACS 2018.[26] The simulations reveal a closer interaction between the negatively charged N-terminal portion of the tail (NCTT) and the positively charged juxtamembrane domain in the active EGFR, while these two domains stay far away from each other in the inactive EGFR (Supplementary Fig. 24) Since the juxtamembrane domain is closer to the plasma membrane, the NCTT is closer to the membrane as well. This is consistent with the results of the coarse-grained simulations reported in the main text.

**Cell Culture and reagents**

CHO cells were purchased from American Type Culture Collection (ATCC) and maintained in 10% $CO_2$ with Ham's F-12K (Kaighn's) medium (Gibco) supplemented with 10% FBS and 1% penicillin-streptomycin (both from Genessee Scientific). Antibodies recognizing the following proteins were purchased: Actin AC-15 (Sigma-Aldrich), EGFR, phospho-EGFR Y1068, phospho-EGFR Y992, phospho-EGFR Y1045, Akt, phospho-Akt S473, p44/42 MAPK (Erk1/2), and phospho-Erk1/2 T202/Y204 (Cell Signaling Technologies). Horseradish peroxidaseconjugated goat anti-mouse and goat anti-rabbit secondary antibodies were purchased from Bio-Rad Laboratories.

**Transfection**

EGFR complementary DNA (cDNA) was sub-cloned into pcDNA3.1(+). The pCDNA3.1+ JM-neu EGFR and CTTneu EGFR plasmids are made by GenScript USA Inc. NJ, USA through mutagenesis. Cells were transfected with polyethylenimine (PEI) reagent with equal amounts of each plasmid. Transfected cells were serum-starved overnight before stimulation with 100 ng/mL epidermal growth factor (EGF) (Millipore-Sigma) for 5, 15 or 30 min in serum-free medium.

**Immunoblotting**

Treated cells were collected and lysed in sample buffer (62.5 mm Tris-HCl, 2% SDS, 5% $\beta$-mercaptoethanol, 10% glycerol, 0.05% bromophenol blue). Lysates were boiled for 5 minutes at 95$^o$ C, resolved by 8% SDS-PAGE, and transferred to nitrocellulose membranes, followed by immunoblotting with the indicated antibodies. Immunoblots were developed using Super-Signal West Femto Maxiumum Sensitivity Substrate (Thermo Scientific) on an Alpha Innotech imaging station. Band density was quantified using ImageJ (National Institutes of Health) and normalized to actin as a loading control.

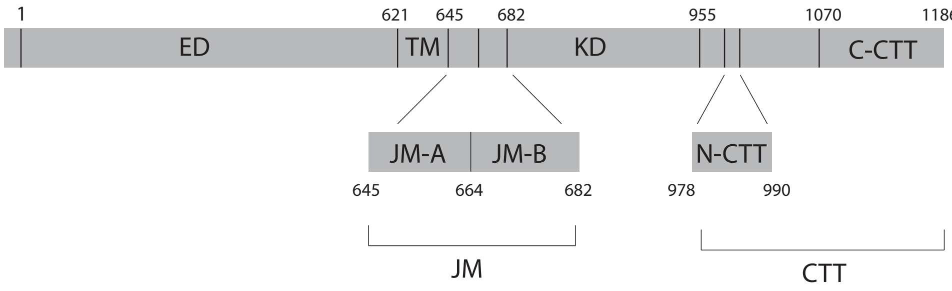


**Supplementary Fig. 1. Domains of EGFR.** EGFR consists of a 621-amino acid extracellular region (ED), a 24-amino acid transmembrane-spanning domain (TM), and an intracellular region, which is a 37-amino acid juxtamembrane domain (JM), a 273-amino acid kinase domain (KD) and a 231-amino acid C-terminal tail (CTT). The JM is further divided into juxtamembrane-A (JM-A) and juxtamembrane-B (JM-B) domains. Residues 978–990 are defined as N-terminal portion of the CTT (N-CTT) and residues 1070–1186 are defined as the C-terminal portion of the CTT (C-CTT). Residue numbering corresponds to EGFR excluding the signal sequence.

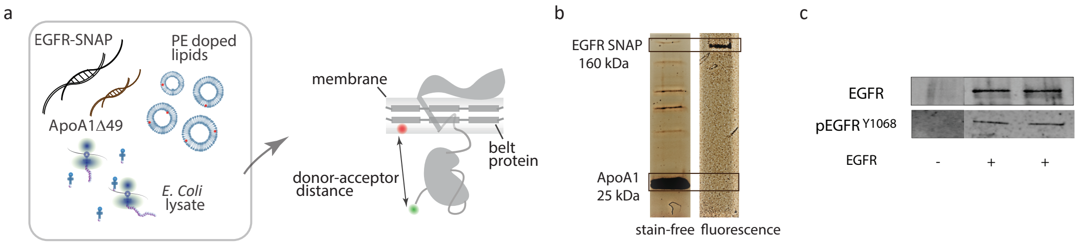


**Supplementary Fig. 2. Production and characterization of full-length EGFR in nanodiscs.** (a) Cell-free reaction for the production of EGFR and nanodisc belt protein. Codon optimized DNA of the receptor protein (EGFR-SNAP) and the belt protein (ApoA1Δ49) are incubated overnight together with lipid vesicles and $E. Coli$ lysate at 25$^o$ C. (b) Stain-free (left) and fluorescence (right) gel images of the His-tag purified sample show the presence of ApoA1 at 25 kDa and full-length EGFR at 160 kDa, which implies successful EGFR production and insertion into nanodiscs. The presence of the EGFR band alone in the fluorescence gel image indicates successful and specific labeling. (c) Western blots were performed on labelled EGFR in nanodiscs. Anti-EGFR Western blots (top) and anti-phosphotyrosine Western blots (bottom) tested the presence of EGFR and its ability to undergo tyrosine phosphorylation, respectively. Bands were observed in both blots when the EGFR plasmid was present, consistent with previous experiments on similar preparations.[1,2]

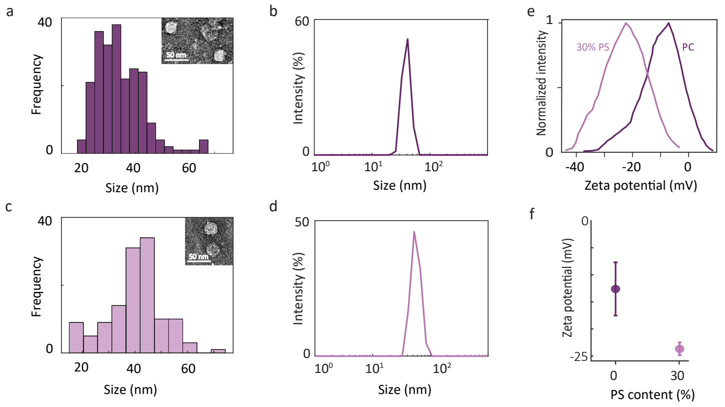


**Supplementary Fig. 3. Characterization of EGFR-containing nanodiscs.** (a) Size distribution of EGFR in DMPC nanodiscs from negative stain transmission electron microscopy (TEM). The circles in the images (inset) show formation of DMPC nanodiscs. The mean disc diameter is 34.1 ± 8.8 nm (N = 224). (b) Dynamic light scattering (DLS) of EGFR in DMPC nanodiscs in PBS buffer indicates ∼37 nm average size, consistent with the TEM results and the FCS curves shown in the Fig. 2g in the main text. (c) Size distribution of EGFR in POPC-POPS (70% POPC/30% POPS) nanodiscs from negative stain TEM. The circles in the images (inset) show formation of POPC-POPS nanodiscs. The mean disc diameter is 40.1 ± 11.0 nm (N = 126). (d) DLS of EGFR in POPC-POPS nanodiscs in PBS buffer indicates ∼53 nm average size, similar to the TEM results. (e) Zeta potential distributions for EGFR in DMPC (dark purple) and POPC-POPS (light purple) nanodiscs. (f) Maximum values from the distributions in (e). Error bars are from five technical replicates.

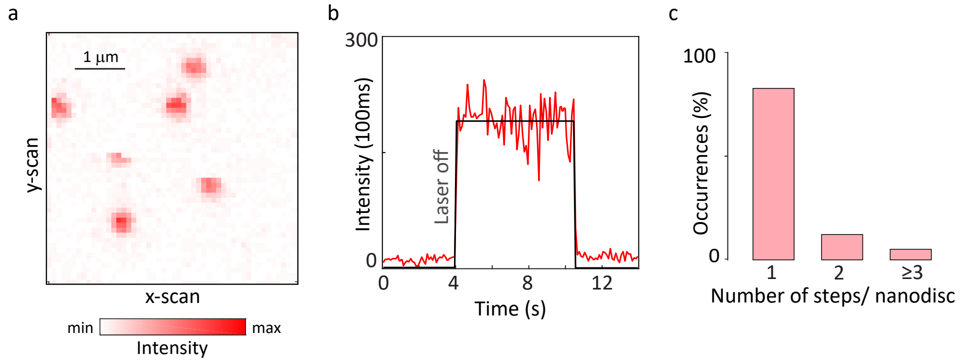


**Supplementary Fig. 4. Photobleaching analysis of Cy5 acceptor molecules in nanodiscs.** (a) Confocal fluorescence image of immobilized constructs of EGFR in nanodiscs containing a labelled Cy5 lipid using 640 nm excitation. (b) Representative intensity time trace from a single construct. The number of detected photons for each 100 ms interval was calculated and used to generate a fluorescence intensity trace (red) with the average intensity for the emissive period overlaid (black). (c) Number of step-wise intensity changes per time trace (N=139). Most molecules (∼85%) exhibited a single-step intensity trajectory indicating the presence of single acceptor molecules.

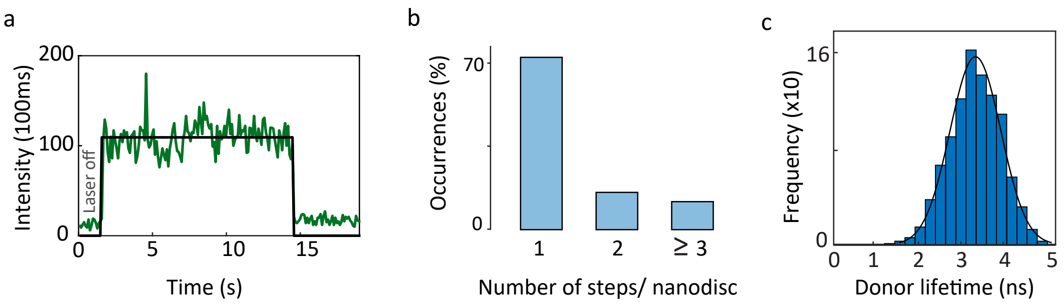


**Supplementary Fig. 5. Photobleaching analysis and lifetime distribution of donor-only constructs.** (a) Representative time trace from single-molecule fluorescence of monomeric donor-only EGFR in nanodiscs showing single-step photobleaching. The number of detected photons for each 100 ms interval was calculated and used to generate a fluorescence intensity trace (green) with the average intensity overlaid (black). (b) Number of step-wise intensity changes per time trace (N=105). The majority of molecules ($\sim$75%) exhibit single-step photobleaching, indicating primarily a single EGFR protein per nanodisc as in previous work.[2] (c) The fluorescence lifetime distribution for the donor-only construct with a median at 3.32 ns.

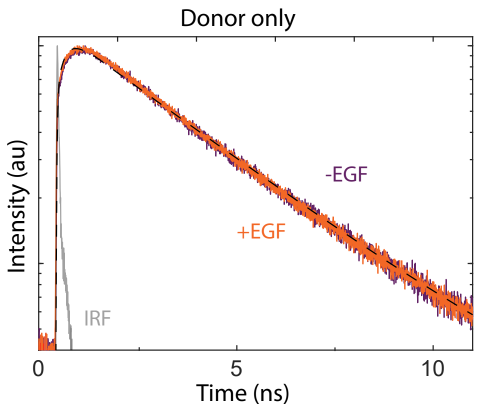


**Supplementary Fig. 6. Ensemble lifetime measurements of snap surface 594.** Ensemble time-correlated single photon counting measurements were performed for the donor dye, snap surface 594 (free dye in 1$\times$ PBS, pH 7.4), in the absence (purple) and presence (orange) of EGF. The instrument response function (IRF) is shown in gray. For both decay traces, a mono-exponential fit (dashed line) found a 3.2 ns lifetime, indicating no photophysical perturbations were induced by EGF.

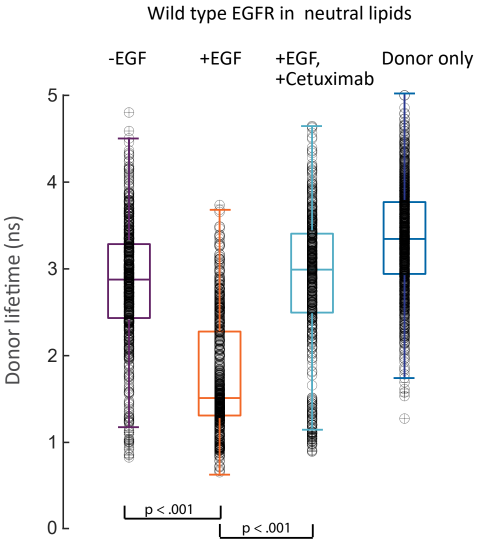


**Supplementary Fig. 7. Box plots of the donor lifetime distributions in a neutral lipid environment.** Distributions of the donor lifetime from the histograms in Fig. 2a of main text are represented as box plots. One-way ANOVA was performed to obtain the P-values (Supplementary Table 6). The donor-only lifetime distribution showed a significant difference compared to other samples (P<0.001). A narrow lifetime distribution with a median of 3.32 ns was found for the donor-only sample. The median value was used as the reference value for calculations of the donor-acceptor distances for all smFRET measurements on WT EGFR in DMPC.

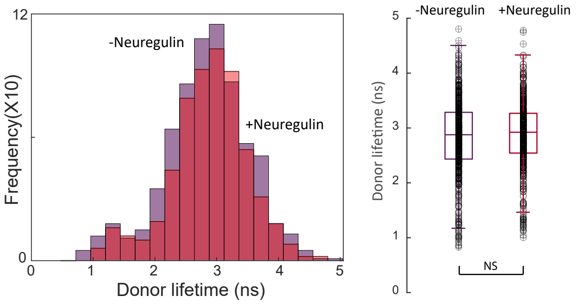


**Supplementary Fig. 8. Lifetime distributions for EGFR in nanodiscs in the presence of neuregulin.** Neuregulin is a structurally similar ligand of the EGF family that does not bind EGFR.[27] smFRET measurements were performed on labelled EGFR in nanodiscs in the presence of neuregulin 1 $\mu$M and the donor lifetime distribution (N=69) was constructed. (left) The distribution is overlaid with the distribution in the absence of ligand . The lifetime distribution in the absence of ligand is taken from main text (-EGF, Fig. 2a, top). (right) The distribution in the presence of neuregulin is statistically similar to the distribution in the absence of ligand (P = 0.2654), indicating the EGF-dependence of the conformational changes reported in the main text. NS, not significant.

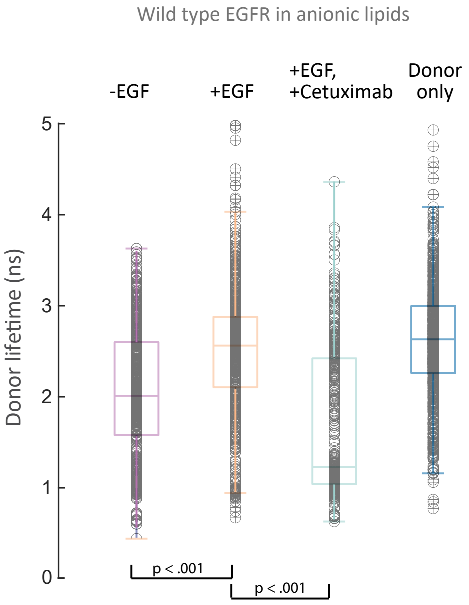


**Supplementary Fig. 9. Box plots of the donor lifetime distributions in a partially anionic lipid environment.** Distributions of donor lifetime from the histograms in Fig. 2b of main text are represented as box plots. One-way ANOVA was performed to obtain the P-values (Supplementary Table 6). The donor-only lifetime distribution showed a significant difference compared to other samples (P<0.001). A narrow lifetime distribution with a median of 2.6 ns was found for the donor-only sample. The median value was used as the reference value for calculations of the donor-acceptor distances for all smFRET measurements on WT EGFR in POPC-POPS.

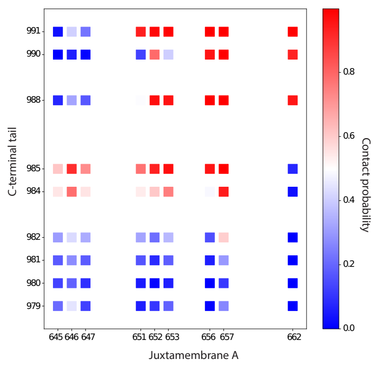


**Supplementary Fig. 10. Critical residues for the electrostatic interaction between the JM domain and the CTT.** The contact probability between the positively charged residues of the JM-A region (residues 645–664) and the negatively-charged residues of the CTT (residues 978–995) was determined using the all-atom simulation of active EGFR. The contact probability was calculated as the fraction of time each pair of residues is closer than 2.0 nm (∼3 times the Debye-Huckel screening length). The residues in the C-terminal half of the CTT have a higher contact probability with the residues in the C-terminal half of the JM-A region (upper right). See text *Section: Atomistic Simulations with the CHARMM Force Field* for simulation details.

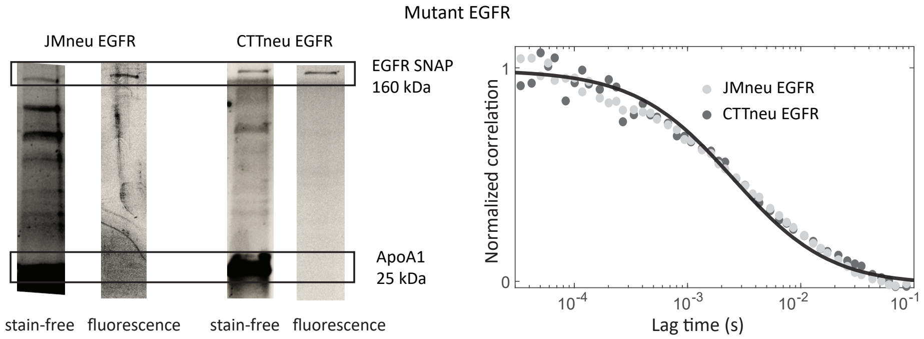


**Supplementary Fig. 11. Characterization of JMneu EGFR and CTTneu EGFR produced *via* cell-free reaction.** (Left) Stain-free and fluorescence gel images show the presence of ApoA1Δ49 at 25 kDa and full-length EGFR at 160 kDa in the His-tag purified sample, which implies successful synthesis and insertion into nanodiscs in cell-free production of JMneu EGFR and CTTneu EGFR. (Right) Fluorescence correlation spectroscopy (FCS) curves of the EGFR mutants in nanodiscs (dots) diffusing through the confocal volume. Solid line represents the fit of the FCS curve for JMneu EGFR. The diffusion timescales (2.54 ms for JMneu EGFR and 2.60 ms for CTTneu EGFR) were comparable to that of WT EGFR (2.2 ms), which shows that constructs of comparable sizes were formed.

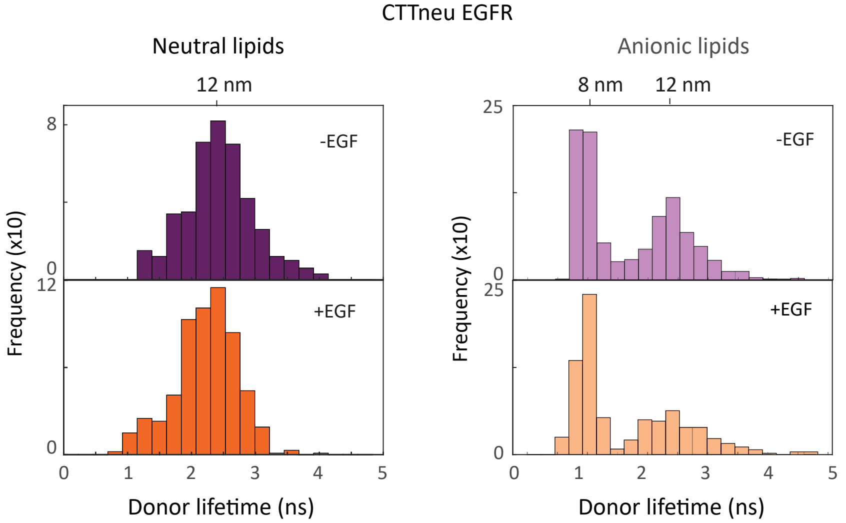


**Supplementary Fig. 12. smFRET measurements of CTTneu EGFR.** smFRET donor fluorescence lifetime histograms for CTTneu EGFR in (left) neutral and (right) partially anionic lipids. Both sets of histograms are statistically similar in the absence (top) and presence (bottom) of EGF (Supplementary Fig. 14), indicating the EGF-dependent intracellular conformational change was lost upon neutralization of the CTT.

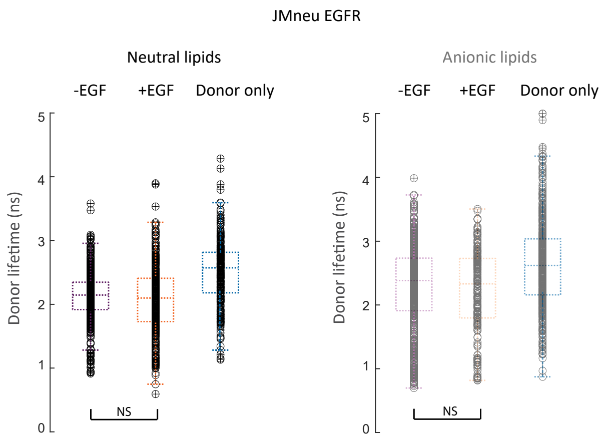


**Supplementary Fig. 13. Box plots of the donor lifetime distributions for the JMneu EGFR mutant in a neutral and partially anionic lipid environment.** Distributions of donor lifetime from the histograms in Fig. 2(e-f) of main text are represented as box plots. One-way ANOVA was performed to obtain the P-values (Supplementary Table 6). The donor-only lifetime distribution showed a significant difference compared to other samples ($P < 0.001$). The donor-only lifetime distribution depends on the local electrostatic environment[28] and so was individually measured for each mutant and for each lipid environment. The medians of the donor-only distributions (2.57 ns in neutral lipids, and 2.61 ns in partially anionic lipids) were used as references to calculate the donor-acceptor distances for the smFRET measurements on JMneu EGFR.

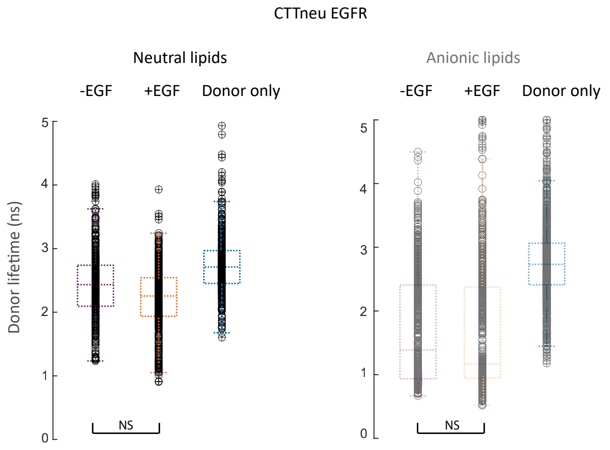


**Supplementary Fig. 14. Box plots of the donor lifetime distributions for the CTTneu EGFR mutant in a neutral and partially anionic lipid environment.** Distributions of donor lifetime from the histograms in supplementary fig. 12 are represented as box plots. One-way ANOVA was performed to obtain the P-values (Supplementary Table 6). The donor-only lifetime distribution showed a significant difference compared to other samples ($P < 0.001$). The donor-only lifetime distribution depends on the local electrostatic environment[28] and so was individually measured for each mutant and for each lipid environment. The medians of the donor-only distributions (2.71 ns in neutral lipids, and 2.73 ns in partially anionic lipids) were used as references to calculate the donor-acceptor distances for the smFRET measurements on CTTneu EGFR.

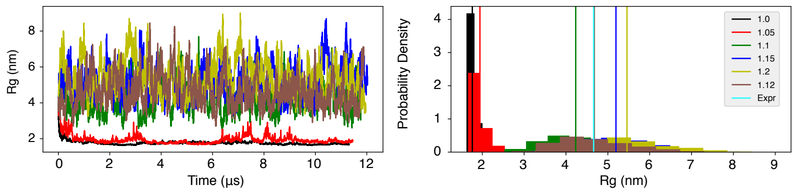


**Supplementary Fig. 15. Adjusting the Martini force field to improve its performance in modeling the CTT portions of the EGFR (residues 961–1186).** The initial configuration of these simulations was prepared with the CHARMM-gui toolkit[23] by solvating the CTT with water molecules and 0.15 M neutralizing ions (NaCl). The CTT structure was built using Modeller.[17] We then carried out independent constant temperature (303K), constant volume simulations with the Lennard-Jones interactions between protein atoms and water molecules scaled by a factor of 1.0, 1.05, 1.1, 1.12, 1.15, and 1.2. Approximately 12 $\mu$s of simulations were performed, and the CTT radius of gyration ($R_g$) as a function of time along different trajectories is shown on the left. The corresponding probability distributions of $Rg$ are provided on the right. The scaling factor of 1.12 (brown) best reproduces the experimental $R_g$ value (cyan) of 4.66 nm. See text *Section: Coarse-grained, Explicit-solvent Simulations with the MARTINI Force Field* for additional simulation details.

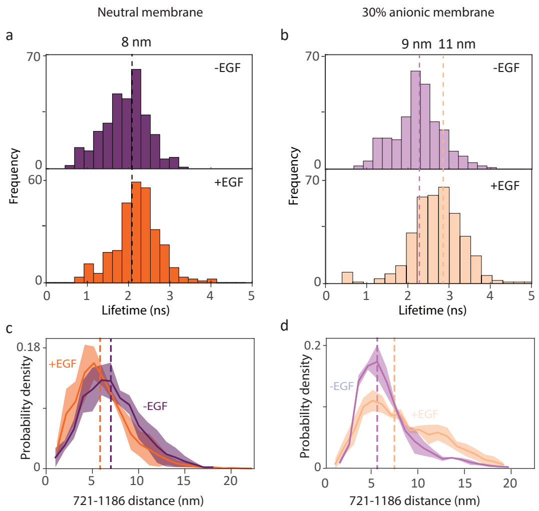


**Supplementary Fig. 16. Comparison of the distance between the ATP binding site and the C-terminus of EGFR from smFRET experiments and simulations.** To further establish the correspondence between the results obtained from experiments and simulations, the distance between the ATP binding site (residue 721) and C-terminus of the protein (residue 1186) was measured using smFRET experiments with a fluorescently-labeled ATP analog and extracted from the simulations for both the neutral and partially anionic lipid bilayers. The results from the smFRET measurements were used to build histograms of the donor lifetime for WT EGFR. (a) In the neutral bilayer, the distribution peaked at ∼8 nm in both the absence (top, purple) and presence (bottom, orange) of EGF. (b) In the partially anionic bilayer, the donor lifetime distributions peaked at 10.9 nm in the presence of EGF and 9 nm in the absence of EGF. The distances between residues 721 and 1186 were calculated from the molecular dynamics trajectories of the simulated conformations in both lipid bilayers and used to generate probability distributions for active (+EGF, orange) and inactive (-EGF, purple) EGFR. (c) In the neutral bilayer, the medians were similar at only ∼1 nm apart whereas (d) in the partially anionic bilayer, they were separated by ∼2 nm, consistent with the experimental trends. The medians for all distributions are given in Supplementary Table 9.

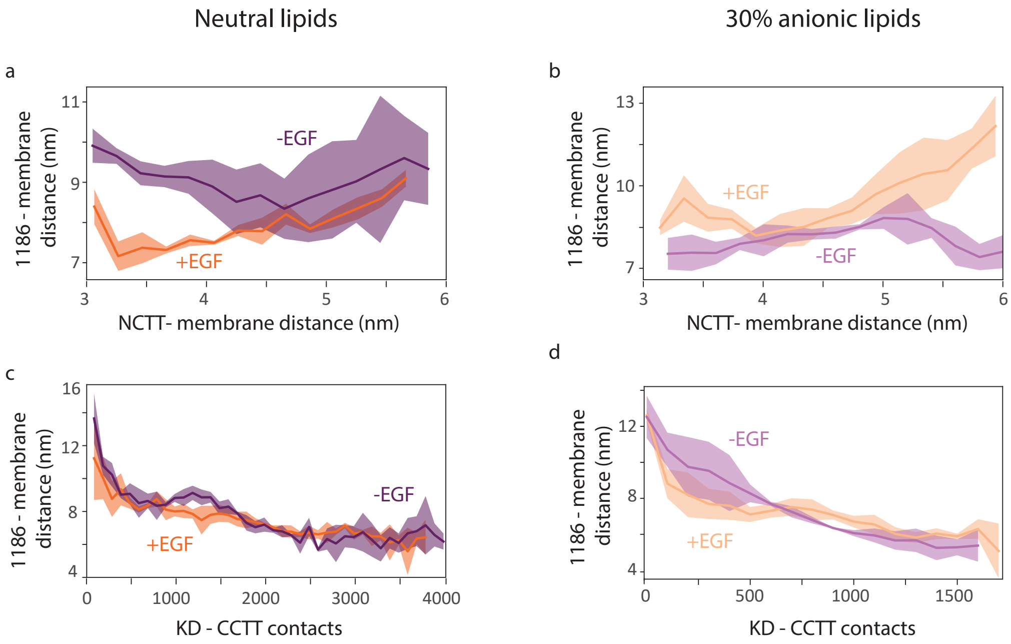


**Supplementary Fig. 17. The distances between the membrane and the C-terminus correlate with two collective conformational variables.** The measured smFRET distance is approximated in the simulations by the distance between the $z$ center of the mass of the lipid bilayer and residue 1186. (a, b) The distance between the bilayer and residue 1186 correlates positively with the distance between the $z$ center of the mass of the lipid bilayer and the N-terminal portion of the tail (NCTT, residues 978–990) for simulations with active (+EGF, orange) and inactive (-EGF, purple) EGFR embedded in neutral (a) or partially negative (b) lipids. (c, d) The distance between the lipid bilayer and residue 1186 correlates negatively with the contacts formed between KD and the C-terminal portion of the tail (CCTT, residues 1070–1186) in the same simulations. Two atoms were recognized as in contact if they were separated by less than 8Å.

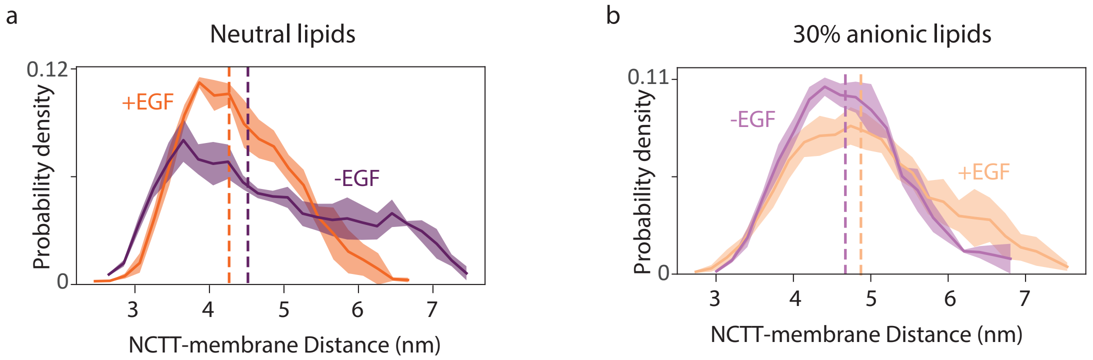


**Supplementary Fig. 18. Probability distributions for the distance between the lipid bilayer and the N-terminal portion of the CTT (NCTT).** The distances between the $z$ center of mass for the lipid bilayer and for the NCTT (residues 978–990) were calculated using simulations with active (+EGF, orange) and inactive (-EGF, purple) EGFR embedded in neutral (a) or partially negative (b) lipids. The median values of the distributions are indicated by dashed vertical lines. The distance reaches the smallest and largest values for the active EGFR in the neutral and anionic membrane, respectively, which reflects the conformational compaction or expansion.

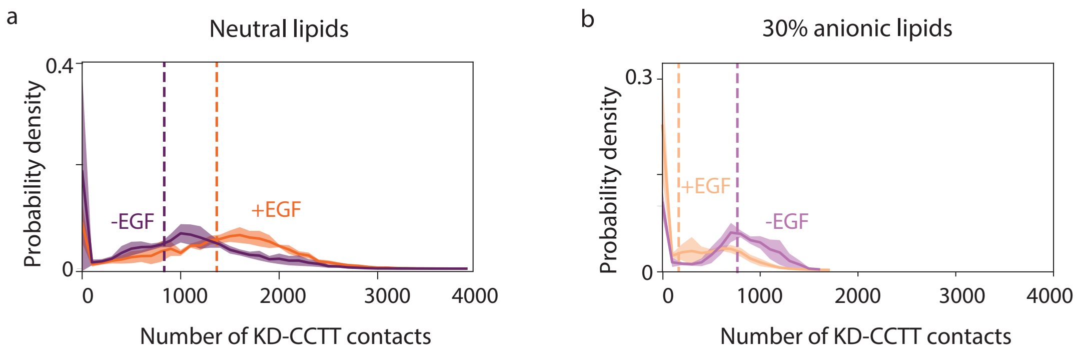


**Supplementary Fig. 19. Probability distributions for the contacts between the KD and the C-terminal portion of the CTT (CCTT).** The number of contacts were calculated between the KD (residues 682–954) and the CCTT (residues 1070–1186) in simulations with active (+EGF, orange) and inactive (-EGF, purple) EGFR embedded in neutral (a) or partially anionic (b) lipids. Two atoms were recognized as in contact if their separation was less than 8Å. The median values of the distributions are indicated by dashed vertical lines. The large number of contacts for active EGFR in the neutral membrane contributes to the conformational compaction.

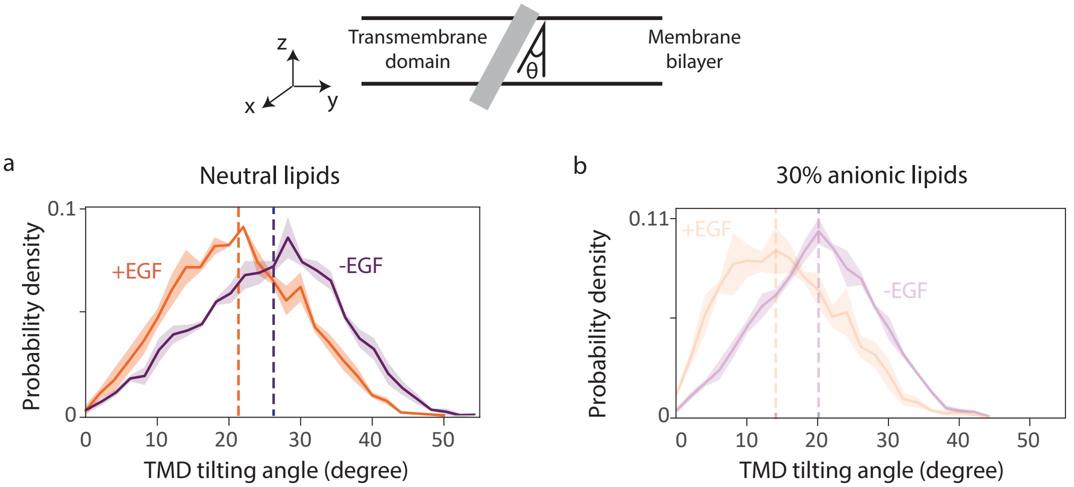


**Supplementary Fig. 20. Probability distributions of the TM domain tilting angle.** The TM domain tilting angle (illustration in top panel) was extracted from simulations with active (+EGF, orange) and inactive (-EGF, purple) EGFR embedded in neutral (a) or partially anionic (b) lipid bilayer. The membrane bilayer stays parallel to the x-y plane throughout the simulations and so the TM domain tiling angle was quantified by measuring the angle $\theta$ between the TM helix and the $z$-axis. The vector along the TM helix was defined using the center of mass coordinates of the N- and C-terminal residues, with residue ID 622 and 644. The median values of the distributions are indicated by dashed vertical lines.

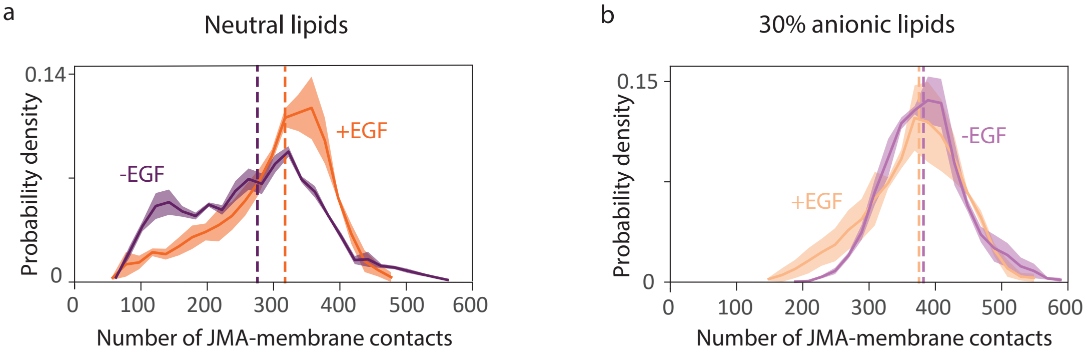


**Supplementary Fig. 21. Probability distributions of the contacts formed between the JM-A region and lipids.** The number of contacts was calculated between the JM-A region (residues 645–664) and the lipid bilayer from simulations of active (+EGF, orange) and inactive (-EGF, purple) EGFR embedded in neutral (a) or partially negative (b) bilayer. Two atoms were defined as in contact if their separation was less than 8Å. The median values of the distributions are indicated by dashed vertical lines. The contacts reflect the degree to which the JM-A embeds into the membrane, which is most significant for inactive EGFR in the anionic membrane.

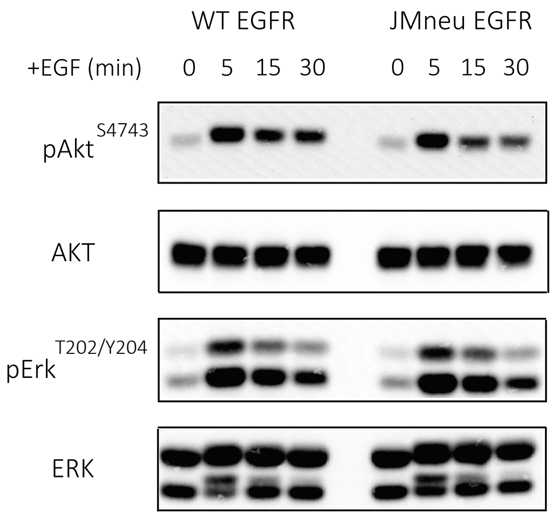


**Supplementary Fig. 22. Cellular experiments with JMneu EGFR show reduced signaling.** The expression level was quantified by Western blot for phosphorylated Akt (pAktS473), phosphorylated Erk (pErkT202/Y204) and total AKT, ERK at 0, 5, 15, and 30 minutes of 100 ng/mL EGF stimulation in CHO cells transfected with WT or JMneu EGFR. Actin was used as a loading control (shown in Fig. 4a in the main text).

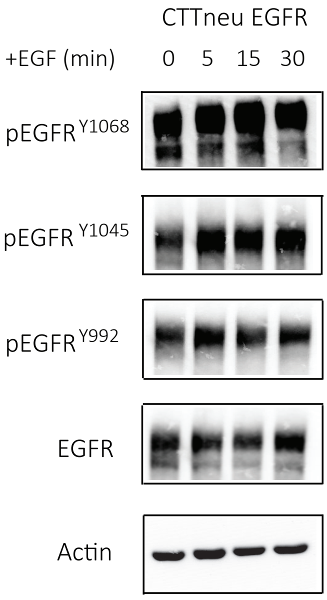


**Supplementary Fig. 23. Cellular experiments with CTTneu EGFR show high basal phosphorylation.** The expression level was quantified by Western blot for phosphorylated EGFR (pEGFRY1068, pEGFRY1045, pEGFRY992) and total EGFR at 0, 5, 15, and 30 minutes of 100 ng/mL EGF stimulation in CHO cells transfected with CTTneu EGFR. Actin was used as a loading control.

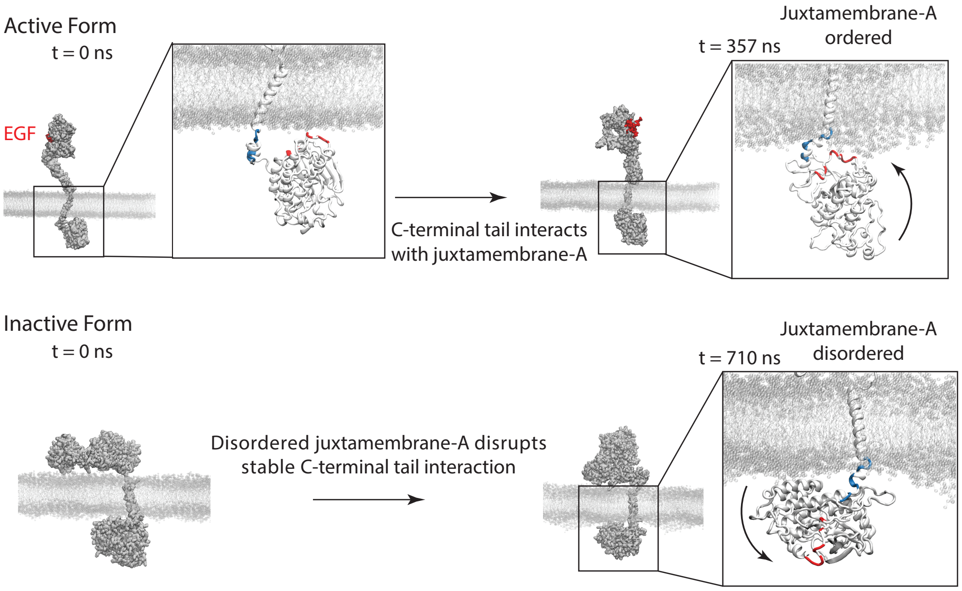


**Supplementary Fig. 24. Representative configurations obtained from all-atom explicit-solvent simulations of the active and inactive (chimera) EGFR with a neutral DMPC bilayer.** See text *Atomistic Simulations with the CHARMM Force Field* for simulation details. Initial configurations of the two simulations constructed from structures reported in Ref.[16] are shown on the left. The juxtamembrane-A (JMA) domain formed close contacts with the N-terminal portion of the EGFR tail (NCTT) as a result of electrostatic interactions in the active simulation, but stayed far apart from each other in the inactive one. The charged residues are colored in blue for positive and red for negative in the insets. Because JMA is located near the membrane, its interaction with NCTT brought the EGFR tail close to the membrane as well. A similar mechanism with smaller NCTT-membrane distance was observed in the MARTINI simulations reported in the main text as well.

| [EGFR] | [Y1068] | Increase in diffusion time | Conclusions |
|---|---|---|---|
| Experiments at 550 nm, monitoring labeled EGFR | | | |
| 1 nM (EGFR nanodiscs) | 0 nM | - | - |
| 1 nM (EGFR nanodiscs) | 0.25 nM | 20.4% | Antibody is bound to pY1068. |
| Experiments at 640 nm, monitoring labelled antibody | | | |
| 0 nM (empty nanodiscs) | 0.25 nM | - | No non-specific interaction of antibody with nanodiscs. |
| 1 nM (EGFR nanodiscs) | 0.25 nM | 3.4% | Antibody is bound to pY1068. |

**Supplementary Table 1. FCS experiments of EGFR phosphorylation.** The Y1068 antibody, which binds specifically to the phosphorylated tyrosine pY1068, was mixed with EGFR labelled with snap surface 594 in nanodiscs. FCS was used to measure the diffusion time of EGFR in nanodiscs $via$ excitation of snap surface 594 ($\lambda_{exc}$=550 nm). The diffusion time increased upon addition of the antibody, consistent with an increase in the hydrodynamic radius of EGFR due to the antibody binding to the phosphotyrosines. FCS was also used to measure the diffusion time of an alexa 647 labelled Y1068 antibody ($\lambda_{exc}$=640 nm). The diffusion time was initially measured in the absence and presence of empty nanodiscs to test for non-specific interactions with the nanodiscs. The same value was extracted (621 $\mu$s) for both measurements. Upon the addition of EGFR nanodiscs, the diffusion time increased (643 $\mu$s), again consistent with binding of the antibody to the phosphotyrosines of EGFR. The binding of Y1068 to labelled EGFR in nanodiscs strongly implies the constructs retain phosphorylation functionality.

| (EGFR:Y1068 antibody) | Donor lifetime (ns) | Decrease in donor lifetime |
|---|---|---|
| 1:0 | 3.08 | - |
| 2:1 | 2.87 | 6.8% |
| 1:2 | 2.68 | 12.9% |
| 1:20 | 1.61 | 47.8% |

**Supplementary Table 2. FRET measurements of EGFR phosphorylation.** Alexa 647 labelled Y1068 antibody, which binds specifically to the phosphorylated tyrosine pY1068, was mixed with EGFR labelled with snap surface 594 in nanodiscs. The lifetime of the donor (snap surface 594) was measured using ensemble time-correlated single photon counting as a function of concentration of the acceptor-labeled (alexa647) Y1068 antibody. The donor lifetime decreased as the antibody concentration increased, which was assigned to an increase in energy transfer from the donor to the acceptor dye. The presence of energy transfer indicated antibody binding to EGFR, because the 8.1 nm Förster radius requires the dye pair be in close proximity. These results are consistent with the FCS measurements (Supplementary Table 1), and also strongly imply the constructs retain phosphorylation functionality.

| Experiments | Experiment distance (nm) | Simulation distance (nm) |
|---|---|---|
| Neutral DMPC membrane | | |
| WT EGFR, -EGF | 11.39 [11.14, 11.65] | 9.22 [8.56, 10.56] |
| WT EGFR, +EGF | 8.14 [8.10, 8.22] | 7.49 [6.83, 7.83] |
| WT EGFR, +Cetuximab, +EGF | 12.02 [11.60, 12.46] | - |
| JMneu EGFR, -EGF | 10.99 [10.82, 11.12] | - |
| JMneu EGFR, +EGF | 10.76 [10.53, 11.06] | - |
| CTTneu EGFR, -EGF | 12.06 [11.58, 12.43] | - |
| CTTneu EGFR, +EGF | 10.97 [10.74, 11.25] | - |
| 30% anionic POPC-POPS membrane | | |
| WT EGFR, -EGF | 10.10 [9.93, 10.33] | 7.38 [6.71, 7.71] |
| WT EGFR, +EGF | *15.22 [13.91, 17.87] | 8.97 [8.30, 10.30] |
| WT EGFR, +Cetuximab, +EGF | 7.98 [7.90, 8.14] | - |
| JMneu EGFR, -EGF | 12.34 [11.90, 12.80] | - |
| JMneu EGFR, +EGF | 11.90 [10.95, 13.12] | - |
| CTTneu EGFR, -EGF | 8.45 [8.17, 9.11] | - |
| CTTneu EGFR, +EGF | 8.00 [7.90, 8.14] | - |

**Supplementary Table 3. Median distances between the membrane and C-terminal end of the protein from smFRET experiments and simulations.** The distance values were extracted from the distributions shown in Figure 2 and Supplementary Figure 12. The numbers in parenthesis indicates the 95% confidence interval for experiments and the minimal and maximal median value from three equal partitions of data for simulations. Asterisk (*) indicates distance was beyond the FRET range for the snap surface 594 and cy5 dye pair (Ro = 8.4 nm).[10]

| Time (minutes) | Reduction in phosphorylation |
|---|---|
| 0 | 56% |
| 5 | 40.1% |
| 15 | 49.4% |
| 30 | 61.9% |

**Supplementary Table 4. Reduction in EGFR phosphorylation of JMneu EGFR.** The level of receptor phosphorylation at Y1068 was quantified from the Western blots shown in Figure 4a. Phosphorylation was reduced for JMneu EGFR compared to WT EGFR in CHO cells.

| Variable | Fit value | 95% confidence interval |
|---|---|---|
| a | 41.8 | [10.9, 72.7] |
| b | 1.6 | [0.3, 2.9] |

**Supplementary Table 5. Intracellular conformational compaction decreased by electrostatic screening.** Parameters from the fit of the percent change in hydrodynamic radius as a function of salt concentration (Fig. 2h in the main text) using Debye-Huckel theory for electrostatic screening.[15] The goodness of fit (R-square) is 0.95.

| Experiment 1 | Experiment 2 | P-value | F | Degrees of freedom |
|---|---|---|---|---|
| DMPC membrane | | | | |
| WT EGFR, -EGF | WT EGFR, +EGF | < 0.001 | 911.3 | 1398 |
| WT EGFR, +EGF | WT EGFR, +EGF, +Cetuximab | < 0.001 | 723.2 | 1254 |
| JMneu EGFR, -EGF | JMneu EGFR, +EGF | 0.0794 | 3.08 | 989 |
| CTTneu EGFR, -EGF | CTTneu EGFR, +EGF | 0.2579 | 1.29 | 139 |
| 30% anionic POPC-POPS membrane | | | | |
| WT EGFR, -EGF | WT EGFR, +EGF | < 0.001 | 172.1 | 1695 |
| WT EGFR, +EGF | WT EGFR, +EGF, +Cetuximab | < 0.001 | 383.1 | 1352 |
| JMneu EGFR, -EGF | JMneu EGFR, +EGF | 0.3636 | 0.83 | 1043 |
| CTTneu EGFR, -EGF | CTTneu EGFR, +EGF | 0.7977 | 0.07 | 1758 |

**Supplementary Table 6. Statistical analysis of smFRET lifetime distributions.** Results from one-way analysis of variance (ANOVA) with P-value, F-statistic and degrees of freedom for all experimental pairs (experiment 1 and experiment 2 in the above table) from the distributions reported in Figure 2 and Supplementary Figure 12.

| Experiments | Number of molecules | Number of bunches |
|---|---|---|
| Neutral DMPC membrane | | |
| WT EGFR, -EGF | 51 | 702 |
| WT EGFR, +EGF | 53 | 697 |
| WT EGFR, +Cetuximab, +EGF | 61 | 558 |
| JMneu EGFR, -EGF | 48 | 563 |
| JMneu EGFR, +EGF | 66 | 427 |
| CTTneu EGFR, -EGF | 59 | 418 |
| CTTneu EGFR, +EGF | 77 | 567 |
| 30% anionic POPC-POPS membrane | | |
| WT EGFR, -EGF | 128 | 891 |
| WT EGFR, +EGF | 121 | 805 |
| WT EGFR, +Cetuximab, +EGF | 65 | 548 |
| JMneu EGFR, -EGF | 96 | 809 |
| JMneu EGFR, +EGF | 66 | 235 |
| CTTneu EGFR, -EGF | 51 | 965 |
| CTTneu EGFR, +EGF | 69 | 794 |

**Supplementary Table 7. Sample sizes for smFRET measurements.** The number of molecules and number of photon bunches used to construct the lifetime distributions are reported for all smFRET histograms from Figure 2 in the main text and Supplementary Figure 12.

| Antibody name | Target | Host | Company | Clone | Dilution |
|---|---|---|---|---|---|
| Anti-EGFR Antibody (A-10) | EGFR C-terminal | Mouse$IgG_{2a}$ | Santa Cruz Biotechnology | Monoclonal | $1:200$ |
| Human Phospho-EGFR Y1068 Antibody | EGFR phosphorylated at Y1068 | Mouse$IgG_{2a}$ | R&D Systems | Monoclonal | $1:200$ |
| Goat anti-Mouse IgG (H+L) Highly Cross-Adsorbed Antibody Alexa Fluor 790 | Mouse | GoatIgG | Thermo Fisher | Polyclonal | $1:10000$ |

**Supplementary Table 8. List of antibodies used to show phosphorylation of EGFR nanodiscs.**

| Experiments | Experiment distance (nm) | Simulation distance (nm) |
|---|---|---|
| Neutral DMPC membrane | | |
| WT EGFR, -EGF | 7.97 [7.83, 8.13] | 7.00 [6.00, 8.00] |
| WT EGFR, +EGF | 8.46 [8.28, 8.57] | 5.84 [5.17, 7.17] |
| 30% anionic POPC-POPS membrane | | |
| WT EGFR, -EGF | 9.03 [8.93, 9.17] | 5.66 [5.66, 5.66] |
| WT EGFR, +EGF | 10.89 [10.50, 11.54] | 7.50 [6.17, 9.17] |

**Supplementary Table 9. Median distances between the ATP binding site (residue 721) and the C-terminal end of the protein (residue 1186) from smFRET experiments and simulations.** The distance values were extracted as the medians of the distributions (Supplementary Fig. 16), which show that the simulations replicate the experimental trends. The numbers in parenthesis indicates the 95% confidence interval for experiments and the minimal and maximal median value from three equal partitions of data for simulations.
